# Characterization of the African Swine Fever Virus transcriptome and the associated innate immune response reveals key features of virulence

**DOI:** 10.1101/2025.10.31.685794

**Authors:** Aurélien Leroy, Juliette Dupré, Chloé Styranec, Pauline Barbarin, Théo Ferré, Vanaïque Guillory, Isabelle Fleurot, Marie-Frédérique Le Potier, Olivier Bourry, Julien Pichon, Sascha Trapp, Grégory Caignard, Ignacio Caballero, Ferdinand Roesch

**Affiliations:** UMR ISP, INRAE, Université de Tours, INRAE Centre Val de Loire, Nouzilly, France; Swine Virology and Immunology Unit, Ploufragan/Plouzané/Niort Laboratory, Agence Nationale de Sécurité Sanitaire de l’Alimentation, de l’Environnement et du Travail (ANSES), Ploufragan, France; Ecole nationale vétérinaire d’Alfort, Anses, INRAE, Laboratoire de Santé Animale, VIROLOGIE, F-94700, Maisons-Alfort, France

## Abstract

African Swine Fever Virus (ASFV) represents a looming threat to animal health, food safety and to the livestock industry. Virulent strains of ASFV cause a severe and often fatal illness, while attenuated strains are usually associated with mild symptoms. Naturally-occurring attenuated strains are typically deleted of more than 20 genes located at the viral genome’s extremities. Whether other key differences between virulent and attenuated ASFV strains may contribute to the virulence phenotype remains however largely unexplored. In this work, we sought to determine how the dynamics of viral gene expression may shape the host’s innate immune response to ASFV infection and contribute to ASFV virulence. We conducted a medium-throughput transcriptomic study to characterize the viral transcriptome of a panel of virulent and attenuated strains (171 viral genes), as well as the host response of ASFV-infected macrophages (92 host genes). Confocal imaging allowed further characterization of cellular response to infection, by assessing the dynamics of IFN and NF-κB pathway activation in ASFV-infected cells.

Our results indicate that the two types of viral pathotypes exhibit global differences in the dynamics of genome replication and viral transcription. Virulent ASFV strains displayed a burst of viral transcription early on, while attenuated strains tended to replicate to higher levels at late time points. The host response was much more pronounced in cells infected with attenuated strains compared to virulent ones, with higher expression levels of interferon-stimulated genes, some innate immunity sensors, and the inducible chaperone HSP70.2. Unexpectedly, genotype I and genotype II virulent strains exhibited some notable differences in their kinetics of viral genome replication and in the host response they provoked, with higher levels of pro-inflammatory cytokines being induced by genotype II strains. Confocal imaging analysis of ASFV-infected primary macrophages revealed that attenuated strains, but not virulent ones, caused the translocation of both p65 and STAT2 to the nucleus. Strikingly, we identified a group of 26 viral genes that were either expressed at higher levels or at an earlier stage of infection by virulent strains. Several of these genes, such as *R298L*, *H233R*, *DP71L* and *MGF505-7R* encode for proteins that inhibited the type I Interferon response in a reporter cell line system. This work sheds new light on the mechanistical drivers of ASFV virulence and will in the long run help to better understand the protection offered by ASFV Live-Attenuated Vaccine candidates.

**Author summary:** African Swine Fever (ASF), a severe infectious disease affecting domestic pigs and wild boars, presents a global threat to the livestock industry. It is caused by African Swine Fever Virus (ASFV), a large DNA virus encoding between 150 and 200 genes. While virulent ASFV strains cause a fatal illness in infected animals, attenuated strains induce only minor symptoms and some can confer subsequent protection against a pathogenic infection. While Live-Attenuated Vaccines for ASFV are under development and represent a promising tool in the fight against ASF, the mechanisms of ASFV virulence (and conversely, attenuation) are not fully understood. In particular, it is unclear whether key differences may exist between attenuated and virulent ASFV strains, beyond the extensive genomic deletions harbored by the former. In this work, we explored for the first time how the dynamics of viral gene expression may influence the innate immune response to different ASFV strains. We found that attenuated ASFV strains trigger a stronger host response compared to virulent ASFV strains, with higher expression levels of innate immune genes and a stronger activation of key signaling hubs. Finally, we identified a group of 26 ASFV genes that may drive this phenomenon and represent novel virulence factors.

## Introduction

African Swine Fever Virus (ASFV) is the causative agent of African Swine Fever (ASF), a high-impact epizootic disease that mainly affects domestic pigs and wild boars. Due to its impact and the absence of widely available medical countermeasures, ASF is listed as a notifiable disease by the World Organization for Animal Health (WOAH). It imposes a considerable socio-economic burden to affected countries, which must implement draconian biosecurity measures and order the culling of infected and at-risk animals to contain outbreaks. Transmission occurs through direct contacts between animals, ingestion of contaminated meat products (swill feeding), and through the persistence of the virus in the environment as fomites (1). ASFV can also be transmitted by bites of *Ornithodoros* soft ticks, and by mechanical transmission by *Stomoxys* stable flies (2,3).

After a first wave of ASF outbreaks in the 1960’s, ASFV was largely eradicated from Europe (4), only to re-emerge in Georgia in 2007. Since then, it has expanded globally, reaching Central and Western Europe; emerging in China in 2018; and circulating rampantly in South East Asia (5). ASF outbreaks can have devastating impacts: in August 2019, over 153 active outbreaks have been reported in China, and some studies have estimated that 300 million pigs were lost from production, representing over 100 billion USD of economic losses (6–8). Although the circulation of the virus has decreased since then, the epidemiological situation is still critical and constantly evolving, as evidenced by recent outbreaks in Germany (9), Italy (10), or Sweden (11).

In its severe forms, ASFV induces a severe hemorrhagic disease, lethal in almost 100% of infected animals in a matter of days (12). In contrast, pigs infected with mildly virulent ASFV strains present only few symptoms from which they quickly recover, and are able to transmit the virus to other animals (13,14). While 24 ASFV genotypes were initially described, a recent re-analysis established 6 main ASFV genotypes (15,16). Genotype I strains were responsible for the ASF epidemic in Europe in the 1960’s, and have resurged in China in 2021 (17). However, the bulk of the current outbreaks is caused by genotype II strains, which all descend from the virulent ASFV strain first isolated in Georgia in 2007 (18). Genotype I attenuated strains include the NH/P68 and OURT88/3 viral isolates, isolated in Portugal from a chronically infected pig and from a soft tick, respectively (19,20). Genotype II attenuated strains have also been described: the Latvia/2017 strain was isolated from a wild boar (21), while the ASFV-989 strain was generated in a laboratory setting, by incomplete thermal inactivation of the virulent Georgia 2007/1 strain (22). Pigs inoculated with such attenuated strains typically develop mild symptoms and are protected against ASFV challenge with closely-related parental strains, making Live-Attenuated Vaccines (LAVs) a promising lead to contain the disease (23). Recently, several gene-deleted vaccine candidates showed good efficacy in protecting against genotype II viruses. For instance, the ASFV-G-ΔI77L vaccine candidate has been licensed and is widely used in Vietnam since 2024. Initial field vaccination results were very promising, with good protection against genotype II strains and a good safety profile (24–27). However, significant concerns emerged quickly: the risk of reversion to virulence (28) or of emergence of vaccine-like strains (29) causing reproductive disorders have been reported. In addition, highly-lethal variants generated by recombination between genotype I and genotype II ASFV strains have been reported and are on the rise in China (30), causing vaccine breakthrough infections (31). Likewise, protection against other heterologous viral strains may also be incomplete (32). Thus, there is still an urgent need to develop innovative prophylactic and therapeutic countermeasures against ASFV and to better understand the mechanisms of protection vaccines may offer.

Experimental infections with virulent and attenuated strains lead to vastly different clinical outcomes, which may be linked to differences in the cytokines produced after infection (33). Indeed, ASFV infects monocytes, dendritic cells and macrophages (34,35), and the cytokine response of these cells to ASFV strains of varying virulence levels seem to differ. *In vitro*, attenuated ASFV strains are associated with an increased interferon (IFN) response compared to virulent ones (36–38). Type I IFN, probably through the action of Interferon-Stimulated Genes (ISGs) like *MXA* or *IFITM3* (39,40), has been reported to have a moderate but significant antiviral effect against ASFV, both *in vitro* and *in vivo* (41,42). This phenomenon may however be strain dependent, with attenuated strains likely being more sensitive to its antiviral effect than virulent ones (37,43). Noteworthy, despite the multiple means for antagonizing the IFN pathway reported (27,44), pigs infected experimentally with highly virulent isolates still produce high levels of IFN (15), which may reflect systemic viral replication and unregulated activation of the immune system. Virulent strains of ASFV also induce a checkless production of pro-inflammatory cytokines (*e.g.* IL6, TNFα, IL1β), the so-called “cytokine storm” (33,45). As is the case with many other viral infections, this can have severe adverse effects on the host, promoting lymphoid depletion, vascular permeability, or provoking hemorrhages in the spleen and other organs (46,47).

The molecular mechanisms driving the establishment of these protective - or at the opposite, deleterious – immune responses remain to be fully characterized. Most of the work on ASFV virulence factors have so far focused on genes deleted in attenuated strains of ASFV: indeed, naturally occurring or laboratory-adapted attenuated strains of ASFV consistently exhibit extensive deletions in the extremities of their genome (48). Deletion of these genes, either individually or in combination, results in a loss of virulence *in vivo*, which is not necessarily correlated with diminished levels of viral replication (49). Various genes considered as ASFV virulence factors were reported to inhibit the IFN response at different levels (44,50), including the cGAS/STING pathway (51), which likely plays a major role in ASFV innate immune detection. Many genes deleted in attenuated strains belong to the Multi Gene Family (MGF) 360 and 505 and have been described to play key roles in the viral interplay with the IFN response (52,53). However, the cellular functions of many putative ASFV virulence factors remain unknown, and the subsets of deleted genes can vary greatly across attenuated isolates. As a result, clearly establishing the determinants of ASFV virulence remains highly challenging.

Multiple transcriptomic studies have used RNA sequencing (RNA-seq) - either bulk sequencing or at the single-cell level - to study the host response to ASFV. Studies conducted *in vivo* showed an important role of host genes like *HTRA3* (54) or of the Netrin pathway (55) in ASF physiopathology. Multiple reports showed significant upregulation of ISGs and of cytokines such as IL18 upon infection (56,57). In contrast, only a handful reports focused on viral transcriptomics. Two studies from Cackett *et al.* investigated the levels of expression of ASFV viral genes in Vero cells infected with the attenuated Ba71V strain (58), or in primary alveolar macrophages infected with the virulent Georgia 2007/1 strain (59). Recently, several single-cell RNA-seq studies quantified host and viral transcription in cells infected with different ASFV isolates (56,60,61). However, the significant cost of these techniques did not always allow to investigate in depth the temporal aspect of this response or to compare a wide panel of ASFV strains of different levels of virulence.

We hypothesized that attenuated ASFV strains, in addition of the deletions they exhibit, may also display different dynamics of transcription in infected cells, which may in turn influence the innate immune response to the virus. Genes encoded by attenuated strains may for instance be expressed at lower levels or later in the viral life cycle than those encoded by virulent strains, and they may represent novel hitherto unknown virulence factors of ASFV. In this work, we used a medium-throughput approach to monitor in detail the kinetics of viral gene expression of a set of two virulent (L60 and OURT88/1) and two attenuated (NH/P68 and OURT88/3) strains, as well as the associated host immune response. We also quantified IFN and NF-κB pathway activation in infected cells over time using confocal microscopy. Our results indicate that the two different types of strains exhibit global differences in the dynamics of genome replication and transcription. We found that 26 viral genes were expressed earlier in the viral life cycle and/or at higher levels in primary alveolar macrophages infected with virulent ASFV strains compared to attenuated ones. In contrast, the host response was clearly more pronounced in cells infected with attenuated strains compared to virulent ones, with higher levels of ISGs; of some innate immunity sensors; and of the chaperone HSP70.2. Further, our results suggest that attenuated strains induce an earlier and more readily detectable nuclear translocation of both p65 and STAT2. These differences in viral genes expression and activation of innate immune pathways could explain the lower pathogenicity of attenuated ASFV strains, and in the long run, may help to better understand the protection offered by Live-Attenuated Vaccines.

## Results

### ASFV strains of varying virulence levels display different replication kinetics

First, we compared the dynamics of viral replication of several genotype I ASFV strains. To this end, primary alveolar macrophages (PAMs) from three different piglet donors were infected with two highly virulent (Lisbon60 and OURT88/1) and two attenuated (NH/P68 and OURT88/3) ASFV strains (**Fig 1A**). To quantify the levels of infection at the end of the first viral life cycle, infected cell cultures were stained at 24 hours post infection (hpi) with antibodies detecting the p72 viral capsid protein and analyzed by flow cytometry. To avoid donor-to-donor variability and to match the infection levels between attenuated and virulent strains, we infected PAMs at two different Multiplicity Of Infection (MOI) with the different ASFV strains. For each viral strain and piglet donor, the MOI was selected to achieve between 20 % and 30 % of p72-positive PAMs (**Fig 1B**). As a result, infection rates were highly similar among piglet donors and ASFV strains conditions (**Fig 1C**). This experimental set up thus allows for robust comparison of other steps of the viral life cycle, such as the levels of viral genomic DNA replication, and viral or host gene expression.

**Figure 1.**
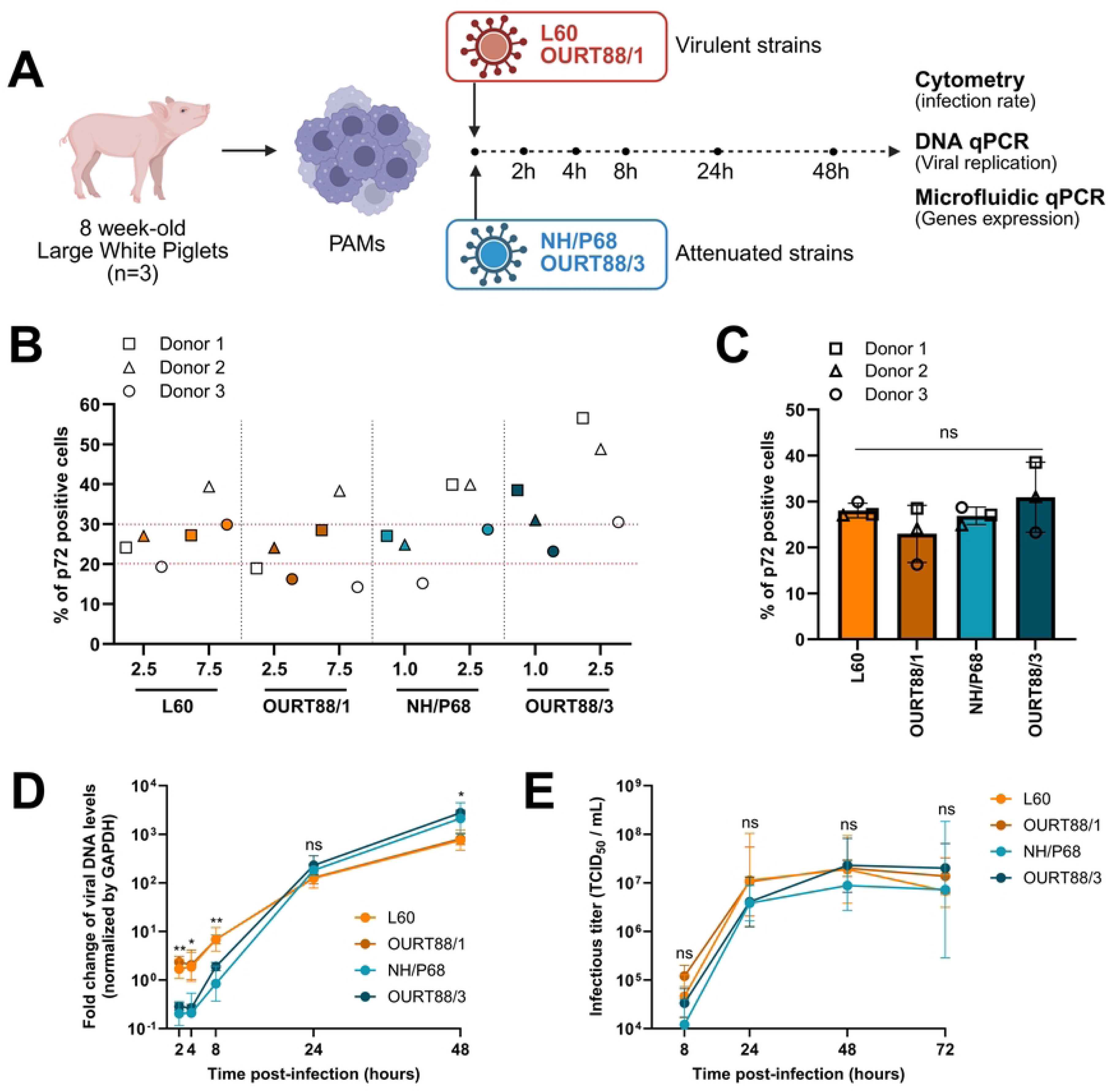
Replication kinetics of virulent and attenuated ASFV strains. **(A)** Experimental workflow. PAMs were isolated from 3 Large White piglets by broncho-alveolar lavage. 5.0 x 10^5^ PAMs were plated in 24-well plates and infected with the indicated ASFV strains. To obtain comparable infection rates at 24 hpi, we adapted the MOI used (1.0 - 2.5 for attenuated strains, 2.5 - 7.5 for virulent strains). Cells were analyzed by flow cytometry at 24 hpi, while DNA and RNA were collected at 0, 2, 4, 8, 24 and 48 hpi for qPCR and transcriptomic analyses. This graphical representation was made using BioRender.com. **(B)** Infection levels at 24 hpi assessed by flow cytometry (all conditions). PAMs isolated from three piglet donors were infected at the indicated MOI with virulent and attenuated ASFV strains. At 24 hpi, infected PAMs were fixed, permeabilized and stained with an anti-p72 antibody. Infection rates were quantified by flow cytometry analysis. The two horizontal doted lines correspond to the 20 % - 30 % range of p72-positive cells. The samples selected for subsequent analysis are colored accordingly to the viral strain used. (**C**) Infection levels at 24 hpi assessed by flow cytometry (selected conditions). Samples with infection rates ranging between 20 % and 30 % were selected and represented. A one-way ANOVA test was performed to compare infection rates among strains (*ns: non-significant*). **(D)** Dynamics of genomic viral DNA replication. Total DNA was extracted at the indicated time points. Viral DNA was quantified by qPCR amplification of the *B646L* gene and normalized against genomic levels of *GAPDH*. An external sample of ASFV-infected PAMs was used for normalization. The geometric mean +/-geometric SD is represented. A two-way ANOVA test was performed to compare viral DNA levels in attenuated or virulent strains infection (*ns: non-significant, *: p<0.05, **: p<0.01*). **(E)** Infectious viral titers. The supernatants of infected PAMs were collected at the indicated time-points, clarified and frozen upon further analysis. Infectious viral titers were quantified by recording the presence of cytopathic effects on PAMs after 7 days. The geometric mean +/-geometric SD is represented. A two-way ANOVA test was performed to compare the production of infectious particles between attenuated and virulent strains infection (*ns: non-significant*).

To quantify the levels of ASFV genome replication, a qPCR assay amplifying the *B646L* gene (encoding the p72 capsid protein) was applied on samples obtained at 2, 4, 8, 24 and 48 hpi. As previously reported (62), ASFV genome replication started between 4 hpi and 8 hpi (**Fig 1D**). Consistently with our flow cytometry data, no significant difference in viral DNA levels was observed at 24 hpi (*p=0.3350*). At 48 hpi, levels of viral DNA were doubled in cells infected with attenuated strains (*p=0.0298*), suggesting an increased replication rate. At earlier timepoints, the levels of viral DNA were ten times higher in cells infected with virulent strains (2 hpi: *p=0.0044*, 4 hpi: *p=0.0332*, 8 hpi: *p=0.0078*). Importantly, these differences in DNA replication levels were not due to the different MOIs used, as PAMs infected at equal MOIs still displayed differences in viral DNA levels (**Fig S1A**). Moreover, even when a higher MOI was used for attenuated strains, PAMs infected with virulent strains still displayed higher levels of viral DNA at early time points (**Fig S1B**). Production of infectious particles over time was assessed by titration of cell culture supernatants at 8, 24, 48 and 72 hpi. Infectious particles were already detected in culture supernatant at 8 hpi, and the peak of viral progeny production (10^7^ TCID_50_ / ml) was reached at 48 hpi for all viral strains. No major difference regarding the production of infectious particles could be detected between attenuated and virulent strains (**Fig 1E**), despite significantly higher levels of viral DNA and transcripts in cells infected with attenuated strains. Taken together, these results suggest that key differences exist in the kinetics of viral life cycle of attenuated and virulent ASFV strains. Attenuated ASFV strains replicate at lower levels at early time points, but at higher levels at late time points. Further, they may display differences in their specific infectivity (*i.e.* the ratio of infectious vs non-infectious particles), since they produce the same amount of infectious viral particles than virulent strains despite higher levels of genome replication at late time points (24 hpi to 48 hpi).

### Dynamics of viral gene expression depend on ASFV virulence

Next, we sought to identify viral genes displaying differences in their temporal expression between attenuated and virulent ASFV strains. We quantified the level of viral gene expression over time for four different ASFV strains using a medium-throughput microfluidic RT-qPCR approach. Specifically, we analyzed the expression of a total of 171 ASFV viral genes, focusing on genes that are conserved across viral strains. More than 90% of the analyzed genes were shared between all strains (158 genes) (**Fig S2A**). Six genes were specific to attenuated strains, while seven were only found in virulent strains. For each viral gene, we determined the timepoint at which its expression became detectable using the microfluidic RT-qPCR assay (**Fig S2B**). Most of the viral genes analyzed were expressed at 8 hpi regardless of the viral strain. Interestingly, we already detected the expression of 70 genes for the virulent strains at 2 hpi, and only 50 genes for the attenuated strains (*p = 0.0001*) (**Fig S2B**). Overall, the profiles and dynamics of viral gene expression were consistent between the two attenuated (NH/P68 and OURT88/3) and the two virulent (L60 and OURT88/1) strains over time (**Fig S2C**). We therefore decided to show the gene expression data in all subsequent analyses as grouped per virulence phenotype, with the two virulent and the two attenuated strains being pooled together.

Consistent with the genome replication data (**Fig 1D**), most ASFV genes were overexpressed at early time points in cells infected by virulent strains compared to those infected by attenuated strains (**Fig 2A-B**). This trend reversed at 24 and 48 hpi, with most viral genes being overexpressed in cells infected by attenuated strains (**Fig 2C-D**). At 4 hpi and 8 hpi, all genes except five (*MGF110-1L*, *MGF110-2L*, *MGF110-9L*, *MGF110-4L* and *86R*) were overexpressed in cells infected by virulent strains. Among these genes, three (*MGF110-2L*, *MGF110*-*9L* and *MGF110-4L*) are specific to attenuated strains, and two (*MGF110-1L* and *86R*) are present in all strains tested. At 24 hpi, only eleven genes remained expressed at higher levels in cells infected with virulent strains, while most other genes were expressed at higher levels in cells infected with attenuated strains (**Fig 2C**). Among them, seven (*MGF360-10L*, *MGF360*-*11L*, *MGF360-12L*, *MGF360-6L*, *MGF505-1R*, *MGF110-12L* and *DP60R*) are deleted from the genome of attenuated strains, and three (*L11L*, *KP86R* and *KP93L*) are present in the genome of both attenuated and virulent strains. The expression kinetics of *L11L*, *KP86R* and *KP93L* genes are presented in supplementary data (**Fig S3A**). At 48 hpi, all ASFV genes (except eight that are deleted in attenuated strains) were overexpressed in cells infected with attenuated strains, and only eight genes were overexpressed in cells infected with virulent strains (**Fig 2D**).

**Figure 2:**
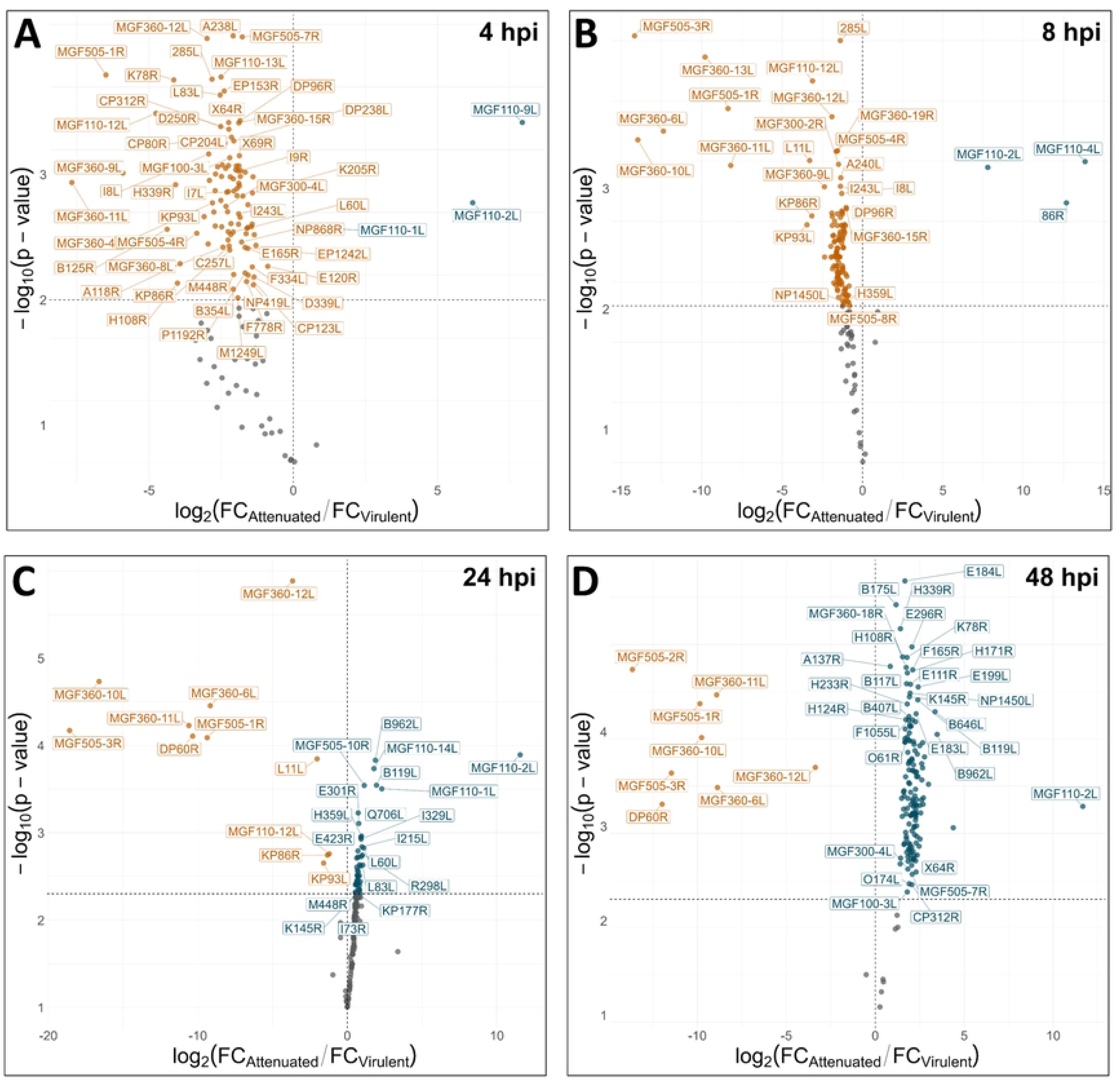
Viral gene expression dynamics show a virulence-specific pattern. **(A-D)** RNA samples were collected at the indicated time points, reverse-transcribed, and cDNA were analyzed by microfluidic qPCR. The expression of each viral gene was normalized against *GAPDH* and *RPL19* expression and fold change values were calculated against the pool of all cDNA analyzed by microfluidic qPCR. Volcano plots representing differentially expressed genes between virulent and attenuated strains at 4 hpi (**A**), 8 hpi (**B**), 24 hpi (**C**) and 48 hpi (**D**) are shown. The X axis represents the log_2_ of the ratio between the mean fold change of attenuated (NH/P68 and OURT88/3) and virulent (L60 and OURT88/1) strains. The Y axis represents the – log_10_ of p-value obtained by an independent t-test comparing the fold change values of virulent and attenuated strains at each time post-infection. Genes with significant upregulation during virulent strain infection are shown in orange (Log_2_FC < 0 and p-value < 0.05), and genes with significant upregulation during attenuated strain infection are shown in blue (Log_2_FC > 0 and p-value < 0.05). Non-significant genes are shown in grey.

Next to the identification of viral genes that are overexpressed by virulent strains or deleted in attenuated strains, these results suggest that ASFV virulence may also be linked to different dynamics of viral gene expression, in particular during the early steps of the viral life cycle.

### Identification of a cluster of 23 genes expressed earlier by virulent ASFV strains

In our analysis, fold change values of viral gene expression closely paralleled DNA levels in infected cells, suggesting that differences in viral abundance between attenuated and virulent strains may have masked subtle changes in gene regulation between strains. To further investigate the kinetics of viral gene expression, we calculated the rate of change in gene expression between consecutive time points for each virus (fold change variation per time increment). This analysis enabled to quantitatively assess and visualize temporal expression dynamics, highlighting genes with the most pronounced transcriptional shifts. Values of derivative fold change were reported on a heatmap where viral genes were clustered accordingly to their variation in expression rates (**Fig 3**).

**Figure 3.**
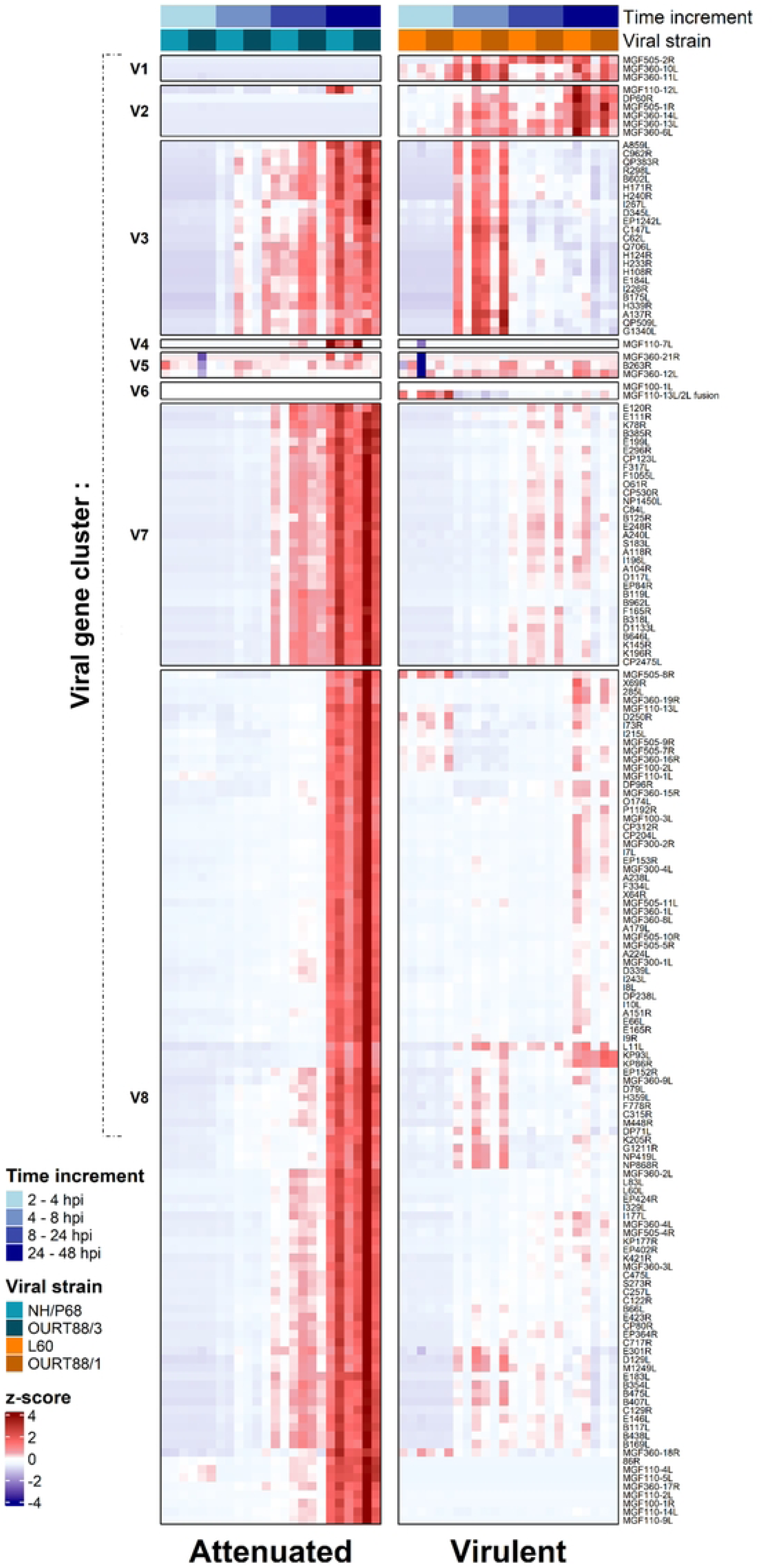
Clustering analysis of viral genes highlights different patterns of gene expression between virulent and attenuated ASFV strains. For each gene monitored by microfluidic RT-qPCR, fold change variation between two consecutive times post-infection was quantified by computing the derivative fold change. In the heatmap, z-score normalized values of the derivative fold change are represented for each gene, each viral strain and at each time increment. Hierarchical clustering based on the derivative fold change data resulted in the classification of viral genes in 8 clusters, annotated V1 to V8. Negative z-score normalized values of derivative fold change (corresponding to the decrease of the fold change) are represented in shades of blue. Positive z-score normalized values of derivative fold change (corresponding to the increase of the fold change) are represented in shades of red.

Hierarchical clustering resulted in the generation of eight viral gene clusters. Clusters V1, V2 and V6 corresponded mostly to Multi Gene Family (MGF) genes that were deleted in attenuated strains. Clusters V7 and V8 grouped the ASFV genes that showed a higher increase in transcription rates at late times post-infection by attenuated strains. These clusters accounted for 75 % of the analyzed genes, and include structural genes (*e.g. CP204L* and *B646L*), MGF genes present in both virulent and attenuated strains, and genes already studied for their role in the disruption of immunity pathways like *A104R* or *I177L*. Cluster V3 grouped 23 genes that were expressed by both the attenuated and virulent strains. This cluster included genes involved in viral transcription, such as *EP1242L* and *C147L*, two subunits of the viral RNA polymerase (63); genes involved in immune escape, like *H240R* and *QP383R* (64,65). The functions of other genes, like *R298L*, *H233R*, or *H171R* were less characterized. In this cluster, the derivative fold change (*i.e.* the rate of increase of gene expression) was higher for the virulent strains compared to attenuated strains during the early times post infection. For virulent strains, it reached its peak between 4 hpi and 8 hpi and leveled off later on. In contrast, for attenuated strains, the biggest changes in expression were observed between 24 hpi and 48 hpi (**Fig 4A**). The detailed expression profiles of cluster V3 genes are available in supplementary data (**Fig S3B**). Interestingly, several genes of cluster V8 (*L11L*, *EP152R, MGF360-9L, D79L*, *H359L, F778R, C315R, M448R, DP71L, K205R, G1211R, NP419L* and *NP868R)* displayed a strong increase in expression between 4 hpi and 8 hpi in cells infected with virulent strains. In contrast, this phenomenon occurred much later in cells infected with attenuated strains (24 hpi-48 hpi). Mapping the different clusters of viral genes on the L60 genome (**Fig 4B**) revealed that many cluster V3 genes are in direct proximity in the genome, with 11 genes being immediately adjacent. Simulations with random shuffling of the cluster classification for L60 genes showed that the spatial proximity of genes belonging to the cluster V3 was not a random event (**Fig 4C**), which may suggest a common transcriptional regulation mechanism shared by these genes. By promoting the early transcription of a set of viral genes, virulent strains may benefit from a “head start” in antagonizing the host response to infection.

**Figure 4:**
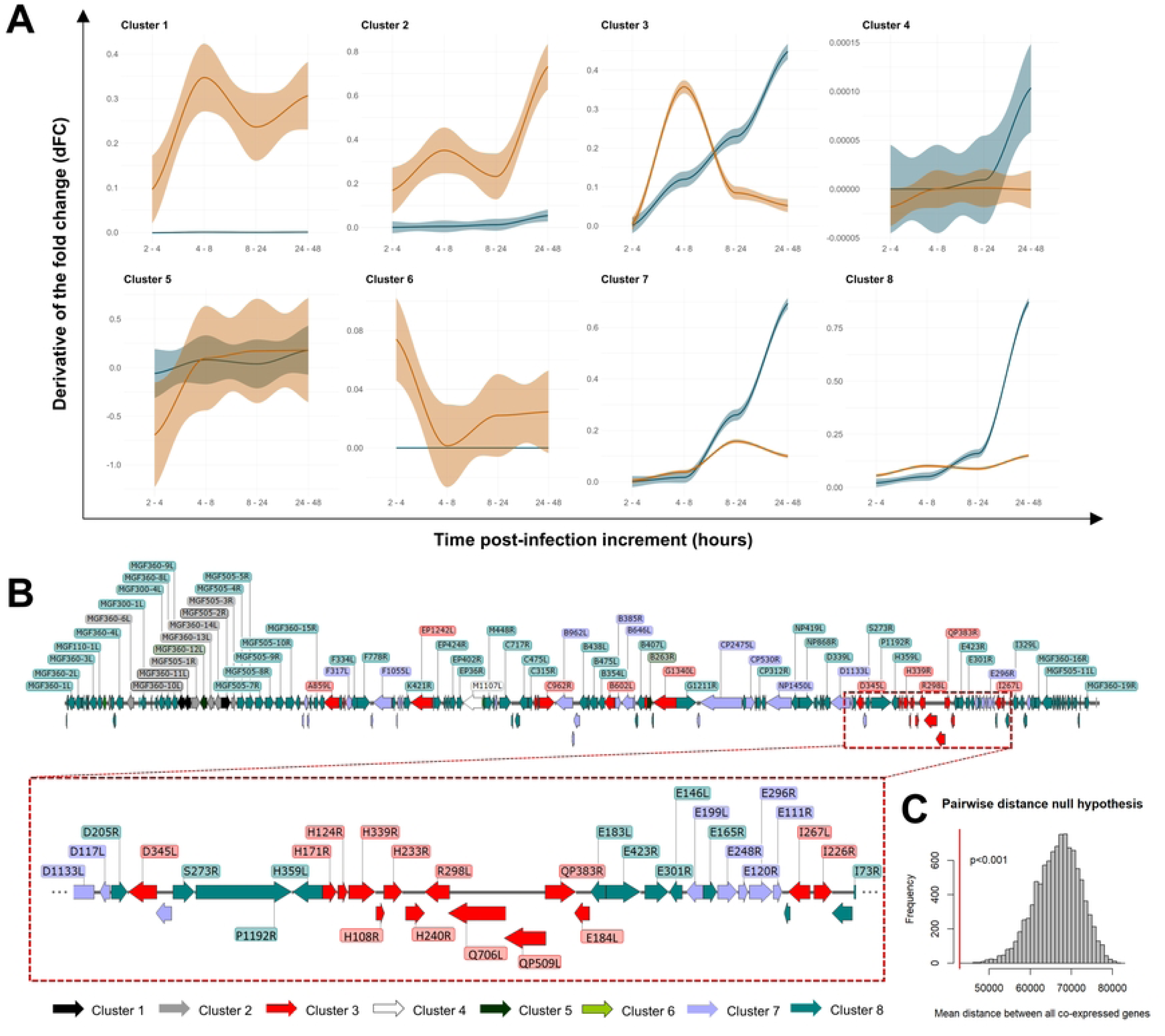
Dynamics of fold change variations over time and mapping of identified viral genes clusters. (A) Dynamics of fold change variations for each viral gene cluster. Trend curves for the 8 identified clusters representing the derivative fold change at each time increment for virulent (orange line) and attenuated (blue line) strain infection. (B) Spatial mapping of viral genes clusters in the L60 genome. The graphical representation of the L60 genome with genes colored according to their classification in clustering analysis was made using SnapGene Viewer software (v6.2.1). (C) Statistical analysis of the distribution of cluster V3 genes in the L60 genome. Histogram showing the distribution of pairwise gene distances obtained after random shuffling of cluster V3 labels. The red line represents the observed value in the original dataset.

### Multiple ASFV genes disrupt the IFN response in a reporter cell line system

Our transcriptional analysis pointed to novel putative virulence factors for ASFV. In order to further characterize their ability to disrupt the host innate immune response, we performed a functional analysis using a luciferase-based IFNβ reporter assay in 293T cells. A total of 14 viral genes were cloned into overexpression plasmid vectors and tested for their ability to interfere with ISRE-driven transcription in response to IFNβ stimulation (**Fig 5A**). Included in this functional analysis were: (i) genes that were overexpressed at 24 hpi during virulent strain infection; (ii) genes that showed a strong transcriptional activity between 4 hpi and 8 hpi (cluster V3); or (iii) those that have a well-documented effect on IFN pathway inhibition.

**Figure 5.**
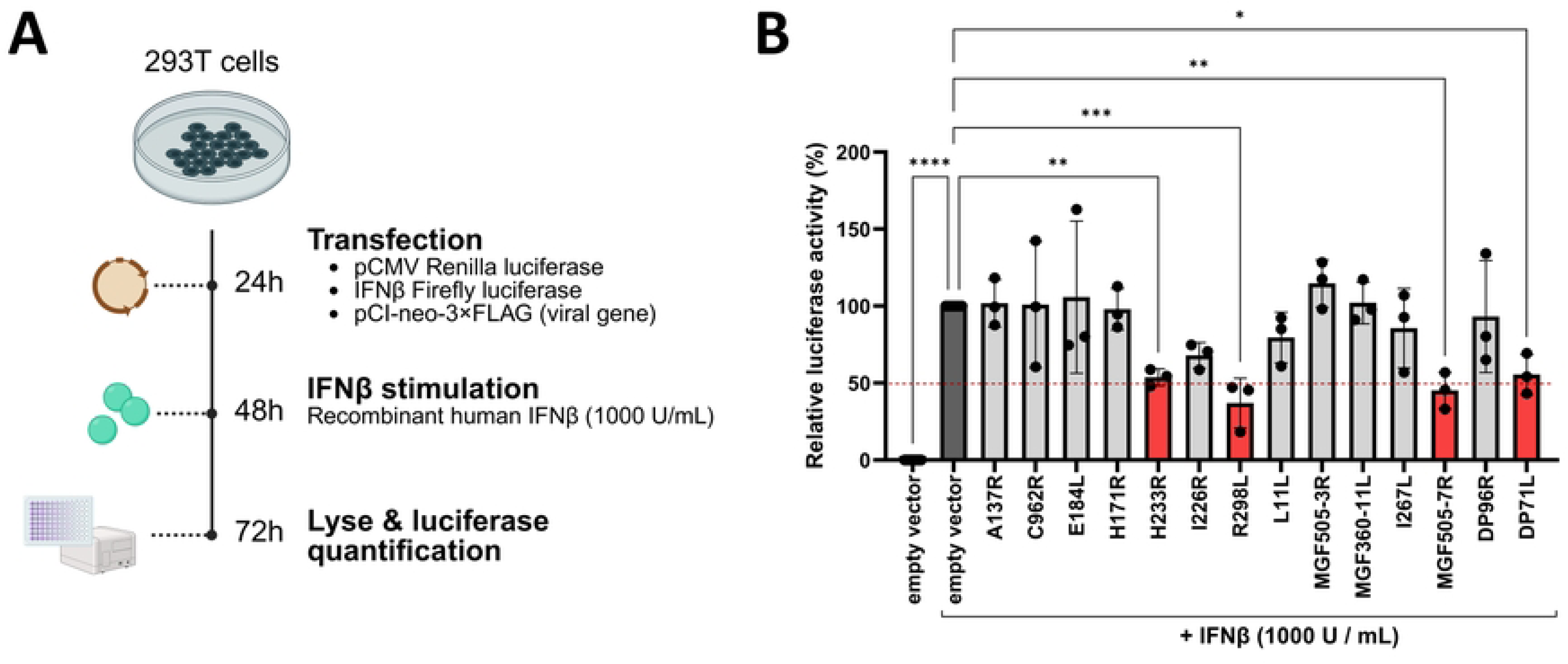
R298L, H233R, DP71L and MGF505-7R inhibit type I IFN signaling in 293T reporter cells. (**A**) Experimental workflow. 293T cells were seeded in 24-well plates at a density of 5.0 x 10^5^ cells per well. 24 h later, cells were transfected with 300 ng of the ISRE-luciferase plasmid, 30 ng of the pCMV renilla luciferase plasmid and 300 ng of the indicated viral gene cloned in the pCI-neo-3×FLAG. 24 h later, cells were stimulated with 1000 U/mL of human recombinant IFNβ. 24 h later, cells were lysed and the relative luciferase activities of IFN-treated cells were determined. (**B**) Inhibition of IFNβ signaling by ASFV viral genes. Firefly luciferase quantification was normalized by renilla luciferase light unit counts to take into account transfection efficiency, cell density and viability in the analyzed samples. The normalized luciferase values were then scaled, fixing the “empty vector + IFNβ” condition at 100 %. Data are represented as the mean +/-SD of three independent experiments performed with 3 technical replicates. Genes with significant downregulation of ISRE activity are shown in red. A one-way ANOVA test was conducted to compare the ISRE activity to the “empty vector + IFNβ” control condition (*: *p* <0.05, **: *p*<0.01, ****: *p*<0.0001).

Both the *R298L* and *H233R* gene products; encoded by neighboring genes within the cluster V3, triggered a significant decrease of ISRE activity upon IFNβ stimulation (60 % and 50 % decrease, respectively) (**Fig 5B**). Interestingly, the protein encoded by the *DP71L* gene, that showed an increased transcription rate during virulent strain infection at early time points, was also able to decrease by 50 % the ISRE activity. No significant effect was detected for proteins encoded by other cluster V3 genes or for *L11L*, which was overexpressed at 24 hpi in cells infected by virulent ASFV strains. Finally, *MGF505-7R* overexpression significantly inhibited IFN signaling, with a 50 % reduction of the IFNβ-mediated luciferase signal. In conclusion, we found that at least four ASFV gene products were able to interfere with type I IFN signaling in our reporter system, those encoded by *R298L*, *H233R, DP71L*, and *MGF505-7R*.

### Differential induction of host pathways in PAMs infected by different ASFV viral strains

Next, we sought to characterize the host response in PAMs infected with different strains of ASFV, using the same microfluidic RT-qPCR approach. The panel of assayed host genes (92 in total) included a number of Interferon-Stimulated Genes (ISGs), as well as genes encoding inflammation mediators, cytokines and chemokines, Pattern Recognition Receptors (PRRs), and surface proteins with putative roles in viral entry. Fold changes in host gene expression were normalized for each piglet donor against the non-infected sample at 0 hpi, and the log2 fold change values were reported on a clustered heatmap (**Fig 6**). These results highlight divergent patterns in the host response to ASFV infection according to the virulence of the strain.

**Figure 6.**
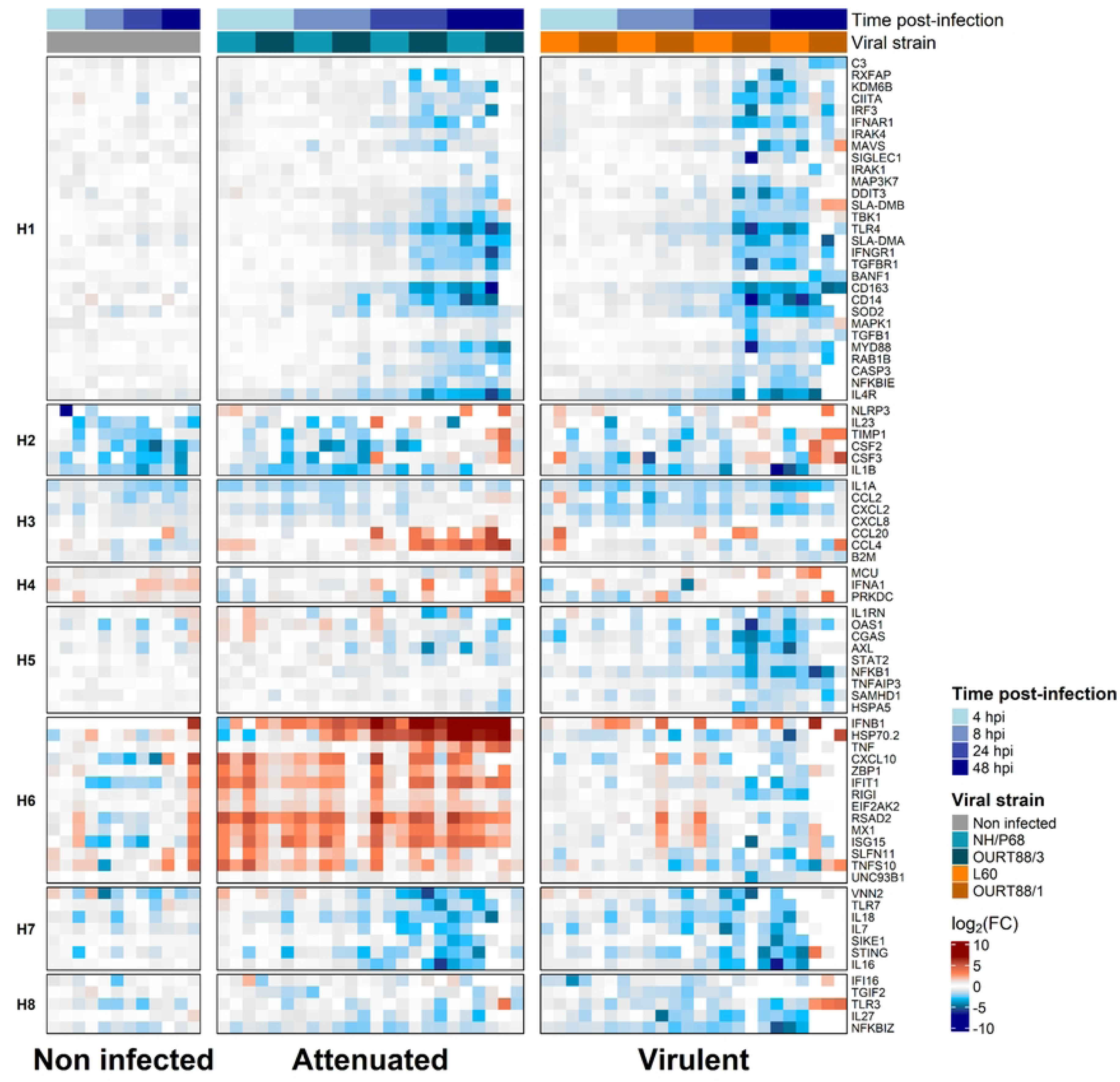
Attenuated strains of ASFV induce a specific host response signature. Heatmap representing the log2 of the fold change of each gene at each time post-infection by each viral strain determined by microfluidic RT-qPCR. For each sample, the expression of each host gene was normalized against *GAPDH* and *RPL19* expression and fold change values were normalized using uninfected PAMs collected at the beginning of the experiment (0 hpi). Hierarchical clustering based on the fold change values resulted in the classification of host genes in 8 clusters, annotated H1 to H8. Upregulated genes are shown in red and downregulated genes in blue. Grey squares represent low or non-varying genes. Undetected genes are shown in white.

First, the expression of housekeeping genes (*RPL19*, *GAPDH*) and of multiple other genes like *B2M* or *TGIF* was similar across infected and non-infected conditions, suggesting that there is no global ASFV-induced shutdown of host gene expression. In contrast, the cluster H6 grouped multiple host genes that were selectively induced in cells infected by attenuated ASFV strains (**Fig 6**), and not in cells infected with virulent ones. For instance, expression of ISGs like *CXCL10*, *ISG15*, *IFIT1* and *MX1* was consistently higher in cells infected with attenuated strains, and was triggered as early as 4 hpi (**Fig 7A; Fig S4A**). Interestingly, a subset of these genes including *IFIT1*, *ISG15* or *EIF2AK2* displayed a clearly discernable second peak of expression at late time points (24 hpi and 48 hpi). *IFNA1* expression was slightly induced by attenuated strains (2-fold at 48 hpi), while *IFNB1* expression was strongly induced at 24 hpi (50-fold) and 48 hpi (>500-fold) by these strains (**Fig 7A; Fig S4A**). In contrast, cells infected with virulent ASFV strains presented no induction of IFN or ISGs at any time point. Infection by attenuated strains also triggered an early upregulation of multiple PRRs: *IFI16* (3-fold), *ZBP1* (6-fold), *EIF2AK2* (2.5-fold), *CGAS* (2-fold) and *RIGI* (3-fold) were all induced at 4 hpi (**Fig 7A; Fig S4A**). Except for *CGAS* and *IFI16*, all these PRRs remained overexpressed in cells infected by attenuated strains at all time points tested. *TNF* expression was also largely increased in PAMs infected with attenuated ASFV, with a fold change over 50 at 48 hpi (**Fig 7A**). Noteworthy, cells infected with attenuated strains displayed elevated levels of *CXCL2*, *CXCL8* and *IL1A* at 48 hpi compared to cells infected by virulent strains (**Fig S4A**). One notable host gene, *HSP70.2*, was massively induced only in cells infected by attenuated strains (∼20-fold at 24 hpi, and >70-fold at 48 hpi) (**Fig 7A**), suggesting the existence of a unique stress response in those cells.

**Figure 7:**
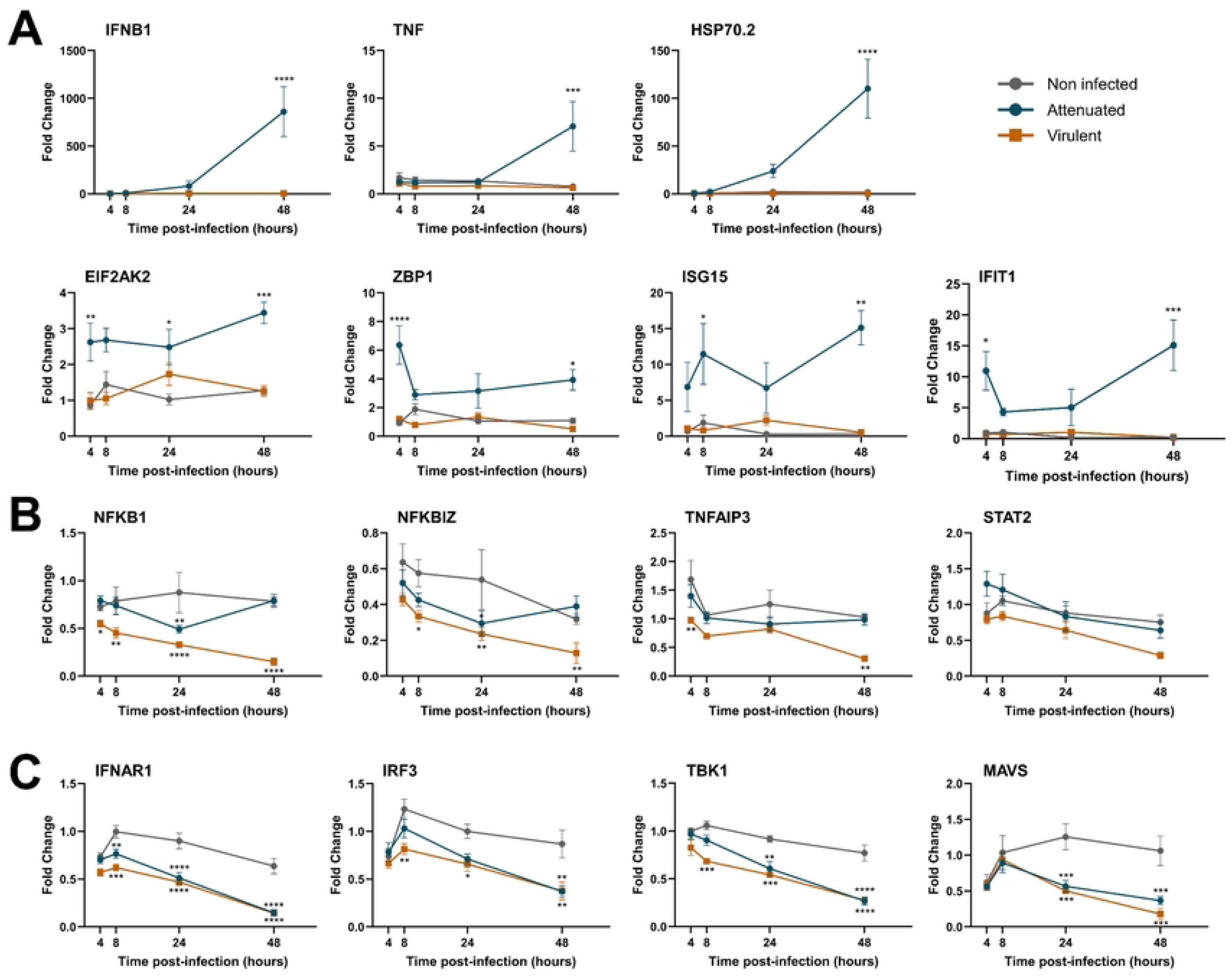
Profiles of expression of host genes in ASFV infected PAMs. (A) Genes specifically triggered by attenuated ASFV strain infection. The microfluidic RT-qPCR data for selected genes of host cluster H6 are shown. (B) Genes downregulated by virulent ASFV strain infection. The microfluidic RT-qPCR data for selected genes of host cluster H5 are shown. (C) Genes downregulated by both attenuated and virulent ASFV strains. The microfluidic RT-qPCR data for selected genes of host cluster H1 are shown. For each sample, the expression of each host gene was normalized against *GAPDH* and *RPL19* expression and fold change values were normalized using uninfected PAMs collected at the beginning of the experiment (0 hpi). A two-way ANOVA test was performed to compare the fold change of gene expression between virulent and attenuated strains infection with non-infected condition at each time-point (*: *p* <0.05, **: *p*<0.01, ***: *p*<0.001, ****: *p*<0.0001).

Expression of host genes from the clusters H5 and H8 (**Fig 6**) was downregulated more efficiently in cells infected by virulent strains. Some of these genes encode proteins related to the NF-*κ*B pathway (**Fig 7B**): at 48 hpi, virulent strains significantly downregulated the expression of *NFKB1* (4-fold), *NFKBIZ* (3-fold) and *TNFAIP3* (3-fold). The downregulation of these genes was already detectable at 4 hpi in cells infected with virulent strains. PAMs infected with virulent strains displayed a modest reduction in *STAT2* levels (2-fold, not significant) compared to uninfected PAMs.

Finally, expression of genes from the clusters H1 and H7 was downregulated by both attenuated and virulent strains, in particular at the late stages of infection (**Fig 6**). These genes mostly encode proteins mostly involved in innate immune signaling; cytokines receptors; as well as surface molecules that may regulate viral entry. The expression of the cognate receptors for IFNα and IFNγ (*IFNΑR1* and *IFNGR1*) was decreased by 2-fold at 24 hpi and 3-fold at 48 hpi (**Fig 7C, Fig S4B**). Similarly, the expression of *TBK1* (3-fold) and *IRF3* (2-fold) was decreased in infected PAMs regardless of the viral strain used. By 48 hpi, the expression of *MAVS* and *MYD88*, key adaptors of the RIG-I and TLR pathways, was decreased by 3-fold. Simultaneously, *IL16* expression was severely impaired by ASFV infection with an 8-fold decrease at 48 hpi. Expression levels of other interleukins such as *IL7*, *IL18* and *IL27* were also negatively impacted. IL4 receptor (*IL4R*) expression was drastically decreased as early as 8 hpi, reaching a 5-fold decrease at 48 hpi. Finally, the expression of genes coding for surface proteins that may participate in viral attachment or entry (such as *CD163*, *SLA-DMA*, *SLA-DMB*, *RXFAP*) were also decreased in the infected PAMs.

Since the expression of PRRs and ISGs was triggered by attenuated strains at 4 hpi, we investigated whether this early induction was a direct consequence of ASFV infection or could represent residual activation of PAMs by the cytokines that may be present in the viral inoculum. Indeed, the ASFV viral stocks are typically generated by infecting PAMs at a low MOI for 5 days. We quantified the levels of porcine IFNα and IFNβ in these viral stocks using a specific ELISA assay. No IFNβ was detected in any of the viral stocks. IFNα was present in the NH/P68 viral production at a low but detectable concentration of 36,72 pg / mL, but not in OURT88/3, the other attenuated strain used (**Fig S5A-B**). While the expression of several ISGs and PRRs was unsurprisingly increased following treatment with recombinant porcine IFNα (**Fig S5C**), this was not the case for HSP70.2 (**Fig S5C**). No major differences were found when we compared the host response to the NH/P68 strain (where residual IFNα could have played a role) and the OURT88/3 strain (in which type I IFN was totally absent) (**Fig S5D-E**). This strongly suggests that carry-over of IFNα in the viral inoculum is not likely to significantly impact our observations.

Overall, both attenuated and virulent strains were able to decrease the transcription of several host genes implicated in innate immune pathways or viral entry. Interestingly, some viral sensors, ISGs, *IFNB1* as well as *HSP70.2* were highly induced by attenuated strains infection but not by virulent strains. These results highlight a virulence-specific pattern in the host response of PAMs infected with different ASFV strains.

### PAMs infected with attenuated ASFV strains display increased levels of p65 and STAT2 nuclear translocation

Next, we aimed to further investigate the activation of NF-*κ*B and IFN pathways triggered by ASFV infection. To ensure optimal detection of pathway activation, PAMs from two piglet donors were infected at a high MOI (MOI 3) with both attenuated (NH/P68, OURT88/3) and virulent (L60 or OURT88/1) ASFV strains. Cells were fixed at multiple time points spanning the first 48 hours of viral infection and analyzed by confocal microscopy. We examined the nuclear translocation of p65, the main subunit of the NF-*κ*B pathway, and STAT2, which translocates in its phosphorylated form upon type I IFN stimulation (**Fig 8**). Representative images of p65 and STAT2 stainings in infected PAMs are presented in **Fig 8A-B**. PAMs were also stained with an anti-p72 antibody detecting the viral capsid in order to monitor infection rates over time (**Fig 8C**).

**Figure 8:**
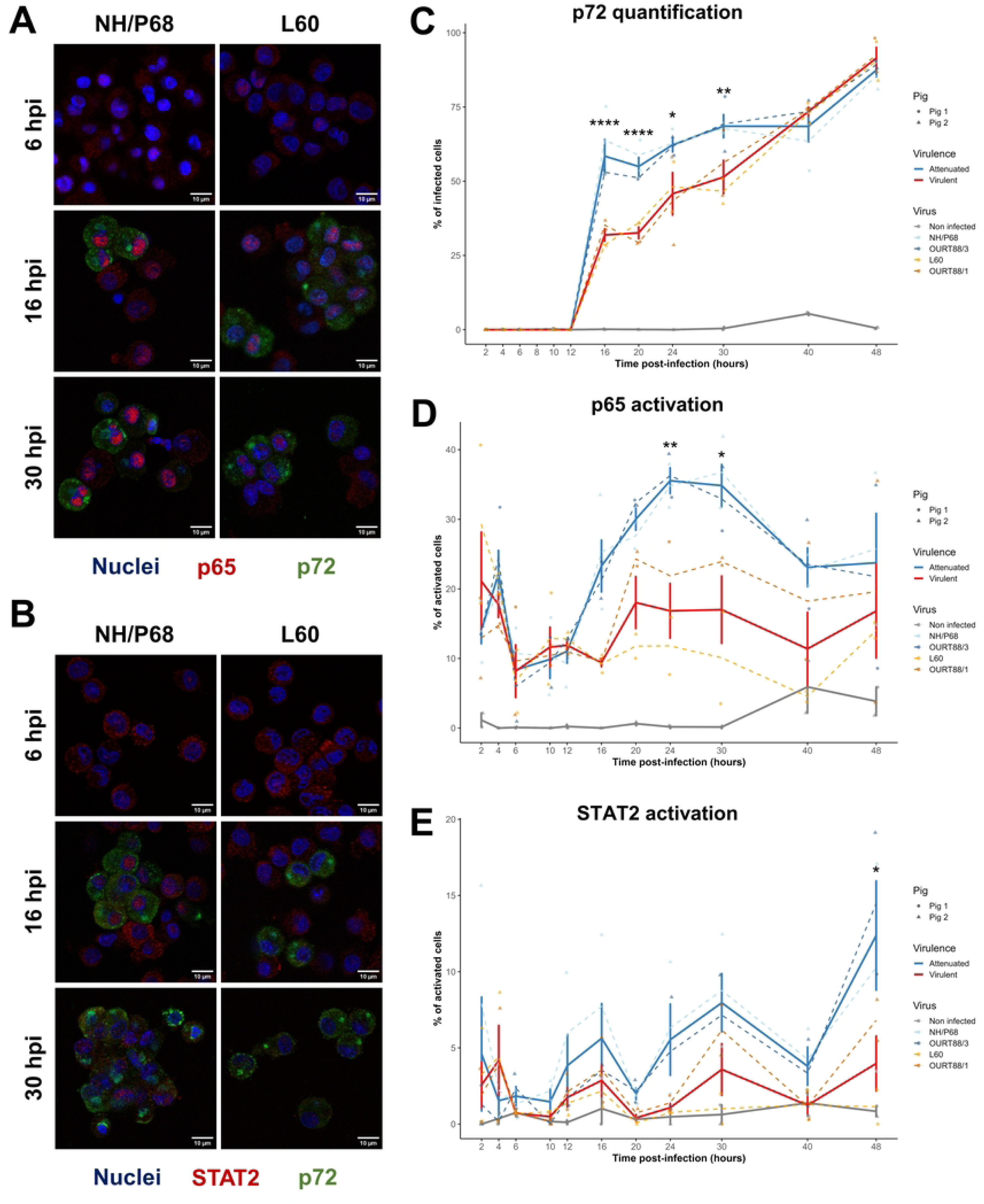
Increased levels of nuclear translocation of p65 and STAT2 in PAMs infected with attenuated strains. **(A**,**B)** Representative confocal microscopy images showing p65 and STAT2 nuclear translocation. 1.5 x 10^5^ PAMs isolated from two piglet donors were plated in 18-well plates suited for confocal microscopy. Cells were infected at an MOI of 3 with the indicated ASFV strains. Representative images at 6, 16 and 30 hpi are shown for p65 (**A**) and STAT2 (**B**) staining. **(C)** Quantification of p72-positive cells over time. A cell is defined as p72-positive when the p72 fluorescence signal is higher than the mean + 3 times the standard deviation of the p72 fluorescence in mock condition (for each pig). A two-way ANOVA test was performed to compare the percentage of p72-positive cells over time (*ns: non-significant, *: p <0.05, **: p<0.01, ****: p<0.001*). (**D**,**E**) Percentage of p65 (**D**) and STAT2 (**E**) activated cells. A pathway in an individual cell is defined as activated when the nuclear-to-cytoplasm ratio of p65 or STAT2 fluorescence is higher than the mean + 3 times the standard deviation of this ratio for the mock condition (for each pig). A two-way ANOVA test was performed to compare the percentage of p65-or STAT2-activated cells over time (*ns: non-significant, *: p <0.05, **: p<0.01*).

PAMs infected with attenuated strains displayed higher rates of p72-positive cells compared to virulent strains from 16 to 30 hpi. For instance, at 24 hpi, 40 % of PAMs were p72-positive for virulent strains, while 60 % of PAMs infected with attenuated strains were p72-positive. At 40 hpi and 48 hpi, the percentage of p72-positive PAMs was similar across conditions. Consistent with the viral DNA replication data (**Fig S1A**), this highlights the ability of attenuated strains to replicate faster than virulent strains starting from the same viral input. The percentage of p65-activated cells was determined by comparing the nuclear-to-cytoplasm ratio of p65 fluorescence between the mock and ASFV-infected conditions over time (**Fig 8D**). Attenuated ASFV strains triggered higher levels of p65 activation in comparison to virulent strains. From 2 hpi to 12 hpi, p65 nuclear translocation was detected in cells infected with both attenuated and virulent strains, with 15-20 % of p65-activated PAMs. A second peak of p65 translocation occurred between 16 hpi and 20 hpi, concomitantly with the first detection of the p72 capsid protein. Between 24 and 30 hpi, p65 nuclear translocation was significantly higher in PAMs infected with attenuated ASFV (30 % of positive cells) compared to PAMs infected with virulent ASFV (20 % of positive cells). While the two attenuated strains tested behaved in a similar way overall, the virulent strain OURT88/1 seemed to trigger slightly more p65 translocation than L60.

Similarly, we evaluated STAT2 nuclear translocation using the nuclear-to-cytoplasm ratio of STAT2 fluorescence (**Fig 8E**). In the first 10 hours of infection, no clear induction of STAT2 translocation was observed in PAMs infected by attenuated or virulent ASFV strains. In both conditions, less than 5 % of the PAMs were considered activated, remaining close to the basal activation state. Starting from 12 hpi, STAT2 seemed to be slightly more translocated in the case of attenuated strain infection compared to virulent strain infection, and this difference became statistically significant at 48 hpi. As noticed for p65, STAT2 seems to also present a cyclic pattern of entries and exits from the nucleus. Oscillation of STAT2 activation seems to increase in strength during attenuated strains infections, whereas the levels of STAT2 activated cells remained almost constant during virulent strains infections. Overall, attenuated strains triggered STAT2 translocation to the nucleus from 16 to 48 hpi while virulent strains did not significantly activate STAT2 translocation at any of the analyzed time points.

### Differences in the host response to genotype I and genotype II ASFV strains

The main experimental approach in this study was to compare virulent and attenuated strains of ASFV. To do this, we used two pairs of genotypes I strains, often used in the literature due to their genetic proximity and similar biological history. Nonetheless, we are also interested in potential differences existing between strains belonging to different genotypes. Hence, we also conducted experiments with PAMs infected by the virulent genotype II strain Georgia 2007/1, responsible for the current panzootic, and compared our results with the other two virulent genotype I strains. These experiments were performed in parallel than those previously described, using PAMs from the same animals.

PAMs infected with ASFV Georgia 2007/1 displayed infection rates comparable to those of genotype I ASFV strains, as assessed by flow cytometry (**Fig 9A**) or by amplification of viral DNA (**Fig 9B**) at 24 hpi. However, the levels of viral DNA in PAMs infected with Georgia 2007/1 was similar to that of the attenuated genotype I strains at all time points evaluated (**Fig 9B**). Notably, levels of viral DNA in PAMs infected with Georgia 2007/1 were significantly lower than in PAMs infected with genotype I virulent ASFV strains at 2 hpi and 4 hpi (**Fig. 9B**).

**Figure 9:**
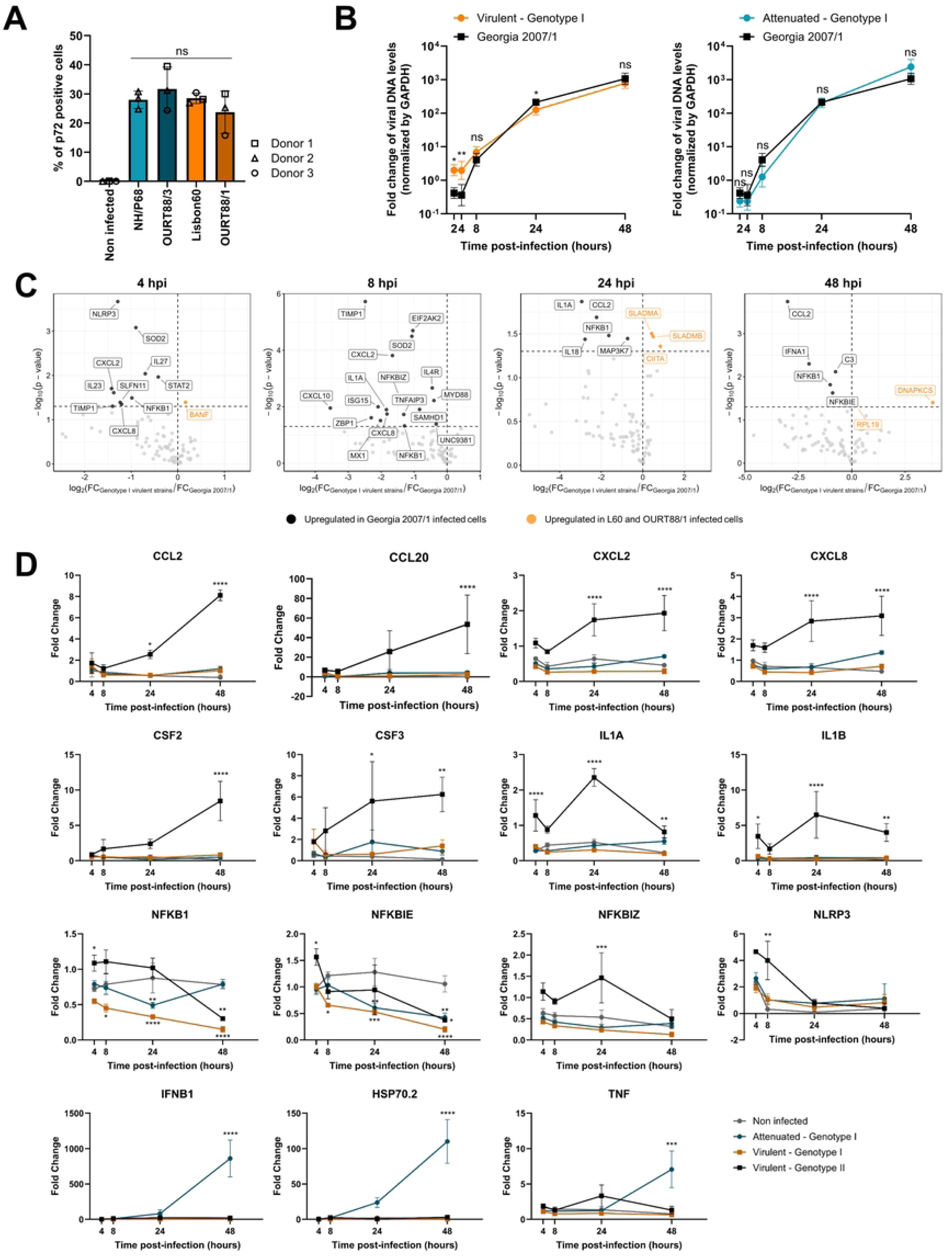
Differences in the host response to genotype I and genotype II ASFV strains. **(A)** Infection levels at 24 hpi assessed by flow cytometry. Infected PAMs were fixed, permeabilized and stained with an anti-p72 antibody. Infection rates were quantified by flow cytometry analysis. A one-way ANOVA test was performed to compare infection rates among strains (*ns: non-significant*). (**B)** Dynamics of viral DNA replication. Total DNA was extracted at the indicated time points. Viral DNA was detected by qPCR amplification of the *B646L* gene and normalized against genomic levels of *GAPDH*. An external sample of ASFV-infected PAMs was used for normalization. The geometric mean +/-geometric SD is represented. A two-way ANOVA test was performed to compare the viral DNA levels in PAMs infected with attenuated genotype I strains versus PAMs infected with Georgia 2007/1 (left panel); or in PAMs infected with virulent genotype I strains versus PAMs infected with Georgia 2007/1 (*ns: non-significant, *: p <0.05, **: p<0.01*). **(C)** Differentially expressed host genes between Georgia 2007/1 and genotype I virulent strains at 4, 8, 24 and 48 hpi determined by microfluidic RT-qPCR and visualized as volcano plots. Each point represents a host gene. For each sample, the expression of each host gene was normalized against *GAPDH* and *RPL19* expression and fold change values were normalized using uninfected PAMs collected at the beginning of the experiment (0 hpi). The X axis represents the log_2_ of the ratio of the mean fold change of genotype I virulent strains (L60 and OURT88/1) and the Georgia 2007/1 strain. The Y axis represents the –log_10_ of p-value obtained by an independent t-test comparing the fold change of virulent strains at each time post-infection. Genes with significant upregulation during Georgia 2007/1 infection are shown in black (Log_2_FC < 0 and p-value < 0.05), and genes with significant upregulation during genotype I virulent strains infection are shown in orange (Log_2_FC > 0 and p-value < 0.05). Non-significant genes are shown in grey. **(D)** Dynamics of expression of host genes induced by ASFV strains infection. The expression data were normalized as in **(C)**. A two-way ANOVA test was performed to compare gene expression among times post-infection of attenuated and virulent strains infection (*ns: non-significant*, *\*: p <0.05*, *\*\*: p<0.01*, *\*\*\*: p<0.001*).

The host response to ASFV Georgia 2007/1 was also quantified by microfluidic RT-qPCR, using cells isolated from the same piglet donors (**Fig 9C-D, Fig S6**). As seen for genotype I virulent strains, ASFV Georgia 2007/1 did not induce the expression of *HSP70.2*, *TNF* or type I interferons (*IFNA1, IFNB1*) (**Fig 9D**). However, it triggered a strong inflammatory response in comparison with the other virulent strains tested. Expression of chemokines *CCL2*, *CCL4*, *CCL20*, *CXCL2* and *CXCL8*, as well as interleukins *IL1A*, *IL1B*, *IL18* and *IL23* and growth factors *CFS2* and *CSF3* were all strongly induced at 24 hpi (**Fig 9D; Fig S6**). The induction of *CCL2*, *CCL20*, *CXCL8*, *CXCL2*, *CSF2* and *CSF3* expression kept increasing over time, reaching a peak at 48 hpi. Between 4 hpi and 24 hpi, Georgia 2007/1 also triggered the expression of genes related to the NF-*κ*B pathway such as *NF*K*B1*, *NF*K*BIE*, *NF*K*BIZ* and *TNFAIP3*, as well as the inflammasome-related gene *NLRP3* (**Fig 9D**). Expression of these genes tend to decrease after 24 hpi. Together, this suggests that genotype I and genotype II virulent ASFV strains may differ in the host response induced in infected PAMs, with a particularly strong induction of inflammatory pathways by genotype II, but not genotype I strains.

## Discussion

Our in-depth analysis of the ASFV transcriptome revealed global differences in the dynamics of the viral life cycle of attenuated and virulent ASFV strains. Infection rates at 24 hpi, as measured by the percentage of cells positive for the p72 antigen (using flow cytometry) or by the amount of viral DNA (using qPCR) were similar for the two types of strains (**Fig 1C-D)**. In contrast, we observed significant differences both in the levels of viral genomic DNA (**Fig 1D**) and viral transcripts (**Fig 2**) during the early (4 hpi – 8 hpi) and late (48 hpi) stages of infection. Indeed, while cells infected with virulent strains tended to express more viral genomes and viral transcripts early on, this trend reversed in the late stages of infection. Consistent with our results, several other studies have reported an increase in viral genomic copies and transcripts in cells infected with attenuated ASFV (59,60), which may be a consequence of the shorter size of their viral genomes. This increase in viral replication observed for attenuated ASFV strains at 48 hpi did not, however, translate in an increase of infectious particles released by infected cells (**Fig 1E**). This raises the interesting possibility that the specific infectivity (*i.e.,* the ratio of infectious vs non-infectious viral particles) of attenuated strains may differ from that of virulent ones. To the best of our knowledge, the increased levels of ASFV DNA and mRNA which we have observed during the first 8 hours of infection by virulent ASFV strains is a novel finding. One interesting hypothesis is that this relates to the differential expression of a specific subset of viral genes, that may favor early genome replication as well as early antagonism of the innate immune response. Strikingly, many of the 23 genes belonging to the viral gene cluster V3, which display a unique profile of transcription dynamics (**Fig 4A**), are localized in close proximity in the viral genome (**Fig 4B**), suggesting a possible co-regulation mechanism. We propose that the genes we identified as being overexpressed at 24 hpi by virulent strains, and those that belong to this cluster of genes which expression is increasing more rapidly for virulent strains, may constitute a group of novel virulence factors that merit to be investigated further. Some of these genes, such as *L11L*, were previously described to be expressed at higher levels by virulent strains, and to impact virulence when deleted (56,66,67). We therefore conclude that viral transcriptomics is a powerful approach to identify novel ASFV virulence factors, which may not be deleted but rather differentially expressed in attenuated ASFV strains.

Using an ISRE-luciferase reporter system, we studied the effect of overexpression of 14 individual ASFV viral genes on IFN signaling (**Fig 5**). We observed a significant decrease in IFNβ-mediated ISRE activity for the proteins encoded by *H233R*, *R298L*, *MGF505-7R* and *DP71L*. While *MGF505-7R* has been extensively studied for its role in ASFV virulence, innate immune escape, and inhibition of the cGAS/STING pathway (68–70), the functions of the other three genes are far less characterized. Two reports suggest that *DP71L*, similarly to *MGF505-7R*, is a virulence factor antagonizing the cGAS/STING pathway (71,72). *R298L* encodes a putative serine/threonine kinase packaged in the viral particle (73). Its deletion from the genome of ASFV Georgia 2007/1 did not seem to impact viral replication or pathogenicity *in vivo* (74). The functions of *H233R* are so far unknown. Our data provide the first functional evidence that H233R and R298L modulate the IFN pathway, making them prime candidates for further study. Importantly, this assay was designed to detect an effect of viral genes in the context of a stimulation of cells by exogenous type I IFN. Therefore, we cannot rule that the genes we tested in this assay display immune escape activity in other context (*e.g.* on the induction of type I IFN synthesis).

Consistent with other reports (38,56,75), we observed that PAMs infected with attenuated strains displayed high expression of *IFNB* and a strong ISG signature, with high expression of *ISG15*, *IFIT1*, *MXA*, among others. Similarly, several Pattern Recognition Receptors (PRRs) like *ZBP1*, *RIGI*, or *CGAS* were also overexpressed at early time points (4 hpi) and maintained a high expression level over time. Although a priming effect of the IFNα (or of other cytokines) present in our viral stocks could not be strictly ruled out, it is unlikely to be solely responsible for this phenomenon, since we obtained similar results when using viral stocks with only small amounts of IFNα (NH/P68) or undetectable traces of IFNα (OURT88/3). In any case, IFN is also produced in significant amounts in experimentally infected animals (38,45) and has a similar effect in this *in vivo* setting. Whether the increased expression of ISGs and PRRs contributes to the overall exacerbated immune response in PAMs infected with attenuated ASFV strains or has a direct antiviral effect remains to be determined. The strong induction of *HSP70.2* in PAMs infected with attenuated strains, also seen in Vero cells infected with the genotype I attenuated strain Ba71V (76), could reflect heightened cellular stress responses. Since **Fig S5C** shows that this phenomenon not driven by IFN, it might represent an interesting IFN-independent defence mechanism.

PAMs infected with attenuated ASFV strains triggered p65 translocation to the nucleus at higher levels than those infected with virulent strains (**Fig 8**), suggesting that the NF-*κ*B pathway is more efficiently activated in those cells. The effect of many virulence factors such as *MGF505-7R*, *A238L* or *MGF360-12L* on NF-*κ*B signaling was previously described (68,77,78) and earlier transcriptomic studies demonstrated differences in NF-*κ*B activation between strains (75). To the best of our knowledge, however, differences in p65 nuclear translocation were not reported before. This early activation may have important downstream effects on the establishment of the innate immune response and/or the synthesis of pro-inflammatory cytokines. Consistent with this idea, we observed a higher induction of pro-inflammatory cytokines like TNFα in cells infected with attenuated ASFV strains (**Fig 7A**). Activation of this pathway by attenuated ASFV strains may have important functional consequences, as it may lead to the induction of pro-inflammatory cytokines such as IL1β and IL6, which have been shown to have a moderate antiviral effect against ASFV (79). In contrast, studies using BAY11-7082, a chemical inhibitor of the NF-*κ*B pathway, and A35, a curcumin-derivative inhibiting NF-*κ*B activation, reported an antiviral effect of these compounds on ASFV replication (80,81). Thus, further work is required to determine whether activation of the NF-*κ*B pathway in infected macrophages has a pro- or antiviral effect on ASFV replication, and whether this effect is dependent on the viral strain tested.

We also investigated the nuclear translocation of STAT2 in cells infected with various ASFV strains. While one study showed increased STAT2 translocation in cells that were stimulated with IFN and subsequently infected with NH/P68 compared to cells that were infected with the virulent Armenia/2008 strain (82), we did not observe such a difference in our work. However, our experimental conditions differed significantly from this study, since we used virulent and attenuated genotypes I strains, and looked at STAT2 localization in absence of any stimulation. Another interesting finding generated by our confocal imaging approach was that p65 and STAT2 activation seems to be biphasic, with a first “wave” of nuclear translocation from 12 hpi to 16 hpi following by another stronger one, between 24 hpi and 30 hpi. Possible explanations for this would be that the first wave of activation corresponds to early viral detection or detecting of incoming virions, while the second is caused by the massive levels of viral replication. Alternatively, ASFV-encoded virulence factors inhibiting p65 and STAT2 translocation may be synthesized at specific steps of the viral life cycle, and/or may no longer fulfill their role when viral replication levels become too important.

The main goal of our study was to compare the expression dynamics and innate immune response induced by genetically-related strains of different levels of virulence. For this, we used two pairs of genotype I ASFV strains which were often compared in the literature, due to their genetic proximity or history of isolation: Lisbon60 and NH/P68, and OURT88/1 and OURT88/3. To add an additional level of comparison, we also used the panzootic genotype II strain Georgia 2007/1. Unexpectedly, this strain did behave differently than the two virulent genotype I strains, in particular regarding the induction of pro-inflammatory cytokines. Indeed, the expression levels of *CCL2*, *CCL20*, *IL1A*, *IL1B* and *TNF* were all more elevated in cells infected with ASFV Georgia 2007/1 compared to those infected with L60 and OURT88/1, two genotype I virulent strains. These results point towards underappreciated differences between virulent strains belonging to different genotypes. Further work is required to determine whether these differences are caused by punctual mutations in key viral genes, or if some viral transcripts are specific to the Georgia 2007/1 strain, as suggested by Cackett *et al* (59). Characterizing the host response triggered by genotype I / genotype II recombinant ASFV strains will also be highly valuable.

Our results shed new light on the mechanisms driving ASFV virulence, which may in the long run contribute to the rational development of ASFV vaccine candidates. A better understanding of the mechanisms of action of ASFV-encoded virulence factors may help to orient the design of gene-deleted LAVs, which ideally need to allow discrimination between vaccinated and infected animals; present good safety profiles; while still retaining enough immunogenicity and sufficient replication levels in infected animals.

## Materials and methods

### Cells and viruses

Primary alveolar macrophages (PAMs) were isolated *post-mortem* from the lungs of 2-month-old Large White (LW) piglets through bronchoalveolar lavage (BAL). Each lung was rinsed using 300 mL of pre-warmed PBS (Sigma-Aldrich, #D8537) supplemented with 1% penicillin/streptomycin (Gibco, #15140122), 1% gentamicin (Euromedex, #EU0540-A), and 1% amphotericin B (MP Biomedicals, #1672348). Collected cells were filtered through a 100 µm cell strainer (BD# 141380C) and washed three times with cold PBS. Red blood cells were lysed using ACK lysing buffer (Gibco #A10492-01). PAMs were then plated in RPMI-1640 medium supplemented with 10% FBS (Gibco, #A5256701) and 1% penicillin/streptomycin (Gibco, #15140122) and cultured at 37°C, 5% CO_2_. Medium was replaced with fresh medium 4 h post-plating. PAMs were typically used in infection experiments 24 h later. In total, three piglet donors provided PAMs for the microfluidic RT-qPCR assay and the tracking of the viral genome replication. PAMs from three more piglets were used for the titration of culture supernatants, and PAMs from two additional piglet donors were used for the confocal kinetics experiment.

293T cell line was purchased from ATCC (#CRL-3216) and cultured at 37°C, 5% CO_2_ in DMEM complemented with 10% FBS (Gibco, #A5256701) and 1% penicillin/streptomycin (Gibco, #15140122).

All viral strains used in this study were provided by the French National Reference Laboratory for African Swine Fever (ANSES). Viral stocks were generated by infecting PAMs at a low MOI (0.1). After 1 h incubation, the media was replaced. Supernatants were collected and clarified when 80% to 90% of cells presented cytopathic effects (typically at 4 – 5 days post-infection). For this study, viruses were passaged a maximum of 5 times on PAMs.

Quantification of viral infectious was performed on PAMs by quantification of cytopathic effects (CPE), as explained in the paragraph below. For the infection experiments, viral stocks were diluted in complete RPMI-1640 medium to the desired multiplicity of infection (MOI) and added to PAMs for 1 h at 37°C, 5% CO_2_. The medium was then replaced with complete medium.

### Titration of culture supernatant

Culture supernatants were clarified by centrifugation at 400 g for 10 min and stored at -80°C until titration. Supernatants were titrated by endpoint dilution in 96-well plates to determine the TCID_50_ titer. PAMs from 3 different piglet donors were seeded in flat bottom 96-well plates at a density of 2.0 x 10^5^ cells per well and inoculated in quadruplet with 10-fold serial dilutions of culture supernatant. To ensure consistency, all samples originating from the same piglet donor were titrated on PAMs isolated from a single animal. Plates were left at 37°C, 5% CO_2_ for 7 days. Wells were then scored for the presence of cytopathic effect, and the TCID_50_ per mL was calculated from the pattern of positive and negative wells using the Spearman-Kärber method.

### Cytometry analysis

PAMs were plated in 12-well plates (1.0 x 10^6^ cells per well) and infected with the different ASFV strains. After 24 h of infection, PAMs were rinsed twice with PBS, detached 30 min at 37°C using Accutase (Millipore #SCR005), and fixed for 15 min at room temperature (RT) using 4% paraformaldehyde (Thermo Scientific #043368.9M). After a step of permeabilization at RT for 20 min in PBS 0.1% BSA 0.2% Triton, cells were stained with the primary anti-p72 mouse antibody (Gold Standard Diagnostics #M.11.PPA.I1BC11) diluted at 1:500 in PBS 0.1% BSA for 1 h at RT. Primary antibody binding was revealed by staining with an anti-mouse Alexa Fluor 647 secondary antibody (Invitrogen #A21236) diluted at 1:500 in PBS 0.1% BSA for 30 min at RT. Flow cytometry analysis was performed using a Fortessa X-20 analyzer (BD).

### Quantification of viral DNA levels by qPCR

DNA of infected PAMs was extracted using NucleoSpin Tissue Kit (Macherey-Nagel #740952.50), following manufacturer instructions. The levels of viral genomic DNA were measured by amplification of the *B646L* gene using real-time qPCR. qPCR was performed using the iQ SYBR Green Supermix (Bio-rad #1708880) and a CFX96 Connect thermocycler (Bio-rad). Each reaction contained 1X iQ Green Supermix, 0.5 µM of forward and reverse primers and 2 ng of the DNA template. The following primers were used: Genomic GAPDH Forward (TCATTTCCTAGGGTGCTG), Genomic GAPDH Reverse (AAGTGTGGTTTTCCCGAGCA), B646L Forward (TGCTCATGGTATCAATATCG), B646L Reverse (CCACTGGGTTGGTATTCCTC). Cycling conditions were as follows: 95°C for 3 min; 40 cycles of 95°C for 10 s and 60°C for 30 s. A melt curve analysis was performed to confirm single-product amplification. *B646L* expression was normalized using *GAPDH* expression and relative quantities were calculated using the ΔΔCt method. Data are presented as geometric mean ± geometric standard deviation (SD) from 3 biological replicates, each measured in triplicates.

### Medium-throughput microfluidic RT-qPCR assay

Total RNA was collected using the Nucleospin RNA kit (Macherey-Nagel #740955.250). An additional DNAse (Macherey-Nagel #740963) treatment of 1 h at 37°C, followed by enzyme inactivation for 10 min at 75°C, was performed directly on eluted RNA to remove viral DNA contamination. cDNA was generated by reverse transcription of 20 ng of RNA using iScript Reverse Transcription Supermix (Bio-rad #1708841), with the following thermal cycle conditions: 5 min at 25°C, 20 min at 46°C and 1 min at 95°C. Pre-amplification was performed with cDNA diluted at 1:10 (for viral genes analysis) or diluted at 1:2 (for host genes analysis) in RNAse free water using a Preamp Master Mix (Fluidigm, #100-5581) in a T100 thermal cycler (BioRad). 1 µl of Preamp Master Mix was mixed with 0.5 µl of a 500 nM primer stock, 2.25 µl of H2O and 1.25 µl of the diluted cDNA (primers sequences are available in **Supplementary Table S1**). Viral primers were initially designed using the L60 genome (GenBank accession number: KM262844.1) and aligned to the genomes of the other viral strains (GenBank accession numbers: NH/P68: KM262845.1, OURT88/3: AM712240.1 and Georgia 2007/1: FR682468.2). In the case of point mutations within the primer sequence, strain/genome-specific primers were designed. Primers targeting host genes were designed using the *Sus Scrofa* genome assembly Sscrofa11.1 (GenBank assembly GCA_000003025.6). All primers were designed using Primer-BLAST (NCBI) with an annealing temperature of 60°C and tested for their amplification efficiency, detection linearity and specificity. The primer sequences are available in Table S1. The pre-amplification thermal cycle conditions were: 95°C for 2 min followed by 14 cycles of 95°C for 15 s and 60°C for 4 min. Pre-amplified cDNAs were subjected to an exonuclease treatment for 30 min (Exonuclease I, New England Biolabs, #MO293) at 37°C followed by 15 min at 80°C and diluted at 1:5 in TE (10 mM Tris-HCI, 1.0 mM disodium EDTA, pH 8.0). Real-time quantitative PCR was performed in a 96x96 Dynamic Array Integrated Fluidic Circuits (Fluidigm), combining 96 samples with 96 primer sets for 9216 simultaneous qPCR reactions. In total, three 96x96 assays were performed to quantify the expression of 171 viral genes and 92 host genes. The reaction mixes for each of the 96 samples was as follows: 3 µl of 2X SsoFast EvaGreen with Low Rox (Biorad, #172-5211), 0.3 µl of 20X DNA binding dye (Fluidigm, #100-7609) and 2.7 µl of pre-amplified cDNA. Specific primer mixes for each of the primer sets were: 3.5 µl of 2X assay loading reagent (Fluidigm, #100-7611), 2.8 µl of TE (10 mM Tris, 0.1 mM EDTA, pH 8.0), 0.35 µl of 100 µM forward primer and 0.35 µl of 100 µM reverse primer. 5µl of the reaction mix and specific primer mix were dispensed into the appropriate inlets and loaded into the circuit. The thermal cycle conditions were 1 min at 95°C, followed by 30 cycles with denaturing for 5 s at 96°C and annealing/elongation for 20 s at 60°C. The quantitative data were analyzed using the ΔΔCt method, where the amount of target was normalized to 2 endogenous reference genes (*GAPDH*, *RPL19*). The results are expressed as fold change (FC) relative to an experimental control consisting of a pool of cDNA from all sample. To calculate the rate of change in gene expression between consecutive time points we approximated the time derivative of the fold change as follows:

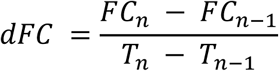

where *FC_n_* is the fold change at time point *T_n_* and *FC_n-1_* is the fold change at the preceding time point *_Tn-1_*. T denotes the time and *dFC* represents the discrete rate of change in fold change between two consecutive measurements (*i.e.,* the derivative fold change).

The full data of viral and host gene analysis are respectively available in Table S2 and Table S3.

### Cloning of ASFV viral genes

All viral gene constructs were cloned by gene synthesis (Twist Bioscience) into pTWIST-ENTR plasmid using the sequences of the Georgia 2007/1 strain. ORFs were then transferred from pTWIST-ENTR into pCI-neo-3×FLAG expression vectors, according to the manufacturer’s recommendations (LR cloning reaction; Invitrogen) and their sequences were verified by sequencing (Eurofins).

### Luciferase Reporter Assays

293T cells were seeded in 24-well plates at a density of 5.0 × 10⁵ cells per well. 24 h later, cells were transfected using jetPRIME reagent (Polyplus) with 300 ng of pISRE-Luc reporter plasmid (Stratagene), which carry the firefly luciferase gene under the control of the ISRE enhancer element. To normalize for transfection efficiency, 30 ng of the pRL-CMV plasmid encoding Renilla luciferase (Promega) was co-transfected in each well, together with 300 ng of either the empty pCI-neo-3×FLAG vector or plasmids expressing the indicated viral genes. Cells were stimulated 24 h later with 1.0 × 10³ IU / ml of recombinant human IFNβ (PBL Assay Science). After 24 h following IFNβ treatment, cells were lysed using Passive Lysis Buffer (Promega). Firefly and Renilla luciferase activities were sequentially quantified using the Bright-Glo and Renilla-Glo luciferase assay systems (Promega), respectively, and luminescence was measured with a GloMax plate reader (Promega). Relative reporter activity was calculated as the ratio of firefly to Renilla luciferase readings. All graphs show mean values and include error bars indicating the standard deviations (SD) of three independent experiments performed in triplicate.

### Confocal microscopy

PAMs collected from 2 different LW piglets were plated in 18-well IBIDI plates (1.5 x 10^5^ cells per well) and infected at MOI 3 with the indicated ASFV strains. PAMs were then fixed using paraformaldehyde 4% for 15 min at 0, 2, 4, 6, 10, 12, 16, 20, 24, 30, 40 and 48 h post-infection. Cells were permeabilized using PBS 1% BSA 0.2% Triton for 30 min at room temperature. Cells were then stained with anti-p72 mouse antibody (Gold Standard Diagnostics #M.11.PPA.I1BC11) diluted at 1:200 in PBS 1% BSA at RT for 1 h. p72-staining was revealed using an anti-mouse Alexa Fluor 488 secondary antibody (Invitrogen #A11029) diluted at 1:500 in PBS 1% BSA at RT for 1 h. Anti-STAT2 and -p65 stainings were performed using rabbit primary antibodies (Cell signaling #72604S, Cell signaling #8242S) diluted at 1:200 in PBS 1% BSA at RT for 1 h. STAT2-and p65-staining were revealed using an anti-mouse Alexa Fluor 594 antibody (Invitrogen #A11012) diluted at 1:500 in PBS 1% BSA at RT for 1 h. Finally, nuclei and actin were stained respectively using DAPI (Sigma #D9542) and Alexa Fluor 647 Phalloidin (Invitrogen #A22287) diluted at 1:10000 in PBS at RT for 20 min. Cells were imaged with a water-immersion 60x/1.2 objective using the Nikon AX R confocal microscope and 405, 488, 560 and 640 nm lasers. Acquisitions were realized using a JOBS protocol. A minimum of 100 cells per piglet donor and per type of strains (virulent or attenuated) was imaged and analyzed. Cell segmentation and fluorescence quantification of p72, p65 and STAT2 proteins in cytoplasm and nucleus of each cell were conducted using QuPath software. Nuclei were segmented using DAPI fluorescence signal and cytoplasm was defined as a 2 µm wide ring surrounding each nucleus. Infection threshold for p72, and activation thresholds for p65 and STAT2, were defined as the mean + 3*SD of the nuclear-to-cytoplasm ratio for each non infected donor. The percentage of infected and activated cells was calculated as the number of infected or activated cells divided by the total number of cells and multiplied by 100. Statistical analysis was performed on RStudio (v4.2.3) and GraphPad10 (v10.5.0).

### ELISA assays

Levels of porcine IFNα and IFNβ in viral stocks have been quantified using commercial ELISA kits (Thermofisher #ES6RB and #ES7RB), following manufacturer instructions. Briefly, 100 µL of viral stocks were diluted 2 times in the kit diluent and incubated in antigen coated wells at RT for 2 h 30 min. After washing, wells were incubated with HRP-conjugated secondary antibody for 45 min. The reaction was developed using TMB substrate, stopped with Stop Solution, and absorbance was measured at 450 nm using a microplate reader (Spark #TCAT93000002).

### PAMs stimulation with recombinant porcine IFNα

PAMs were collected as described earlier in this section, and plated in 24-well plates at a concentration of 5.0 x 10^5^ cells per well. 24 h later, PAMs were stimulated with a low (1 U / mL) or a high dose (250 U / mL) of recombinant porcine IFNα (R&D Systems, #17105-1) diluted in 500 µL of RPMI-1640 medium supplemented with 10% FBS (Gibco, #A5256701) and 1% penicillin/streptomycin (Gibco, #15140122). After 24 h of stimulation, RNA samples of stimulated PAMs were extracted and stored at -80°C until analysis in RT-qPCR. Gene expression was quantified by real-time qPCR using iQ SYBR Green Supermix (Bio-rad #1708880) on a CFX96 Connect thermocycler. cDNA was synthetized from 100 ng of total RNA using iScript Reverse Transcription Supermix (Bio-rad #1708841), following manufacturer instructions. Reaction contained 1X iQ Green Supermix, 0.5 µM of forward and reverse primers and 2 ng of cDNA template. Cycling conditions were: 95°C for 3 min; 40 cycles of 95°C for 10 s and 60°C for 30 s; followed by a melt curve analysis to confirm single-product amplification. The expression of target gene s was normalized to *GAPDH* expression and relative quantities were calculated using the ΔΔCt method. Data are presented as mean ± SD from 3 biological replicates, each measured in duplicates.

### Statistical analysis & graphical representations

To assess the genomic distance of cluster v3 genes, we quantified spatial proximity by calculating the mean of all pairwise distances between their genomic start coordinates. Statistical significance was evaluated through 10,000 label permutations to generate a null distribution. Empirical p-values were derived as the proportion of permutations with mean distances less than or equal to the observed value.

The ANOVA tests were performed on GraphPad Prism (v10.5.0) and are indicated in each figure legend. The t-tests presented in volcano plots were performed on RStudio using *rstatix* package.

The Venn diagram presented in **Fig S2A** was generated with R software (v.4.2.3) using *ggvenn* package. The principal component analysis (PCA) presented in **Fig S2C** was performed on R software using the *FactoMineR* package and visualized using the *factoextra* package. Prior to PCA, the fold change of all genes shared by both virulent and attenuated strains were normalized and scaled (mean centered and divided by the standard deviation). The heatmap representing the derivative fold change of the viral gene expression at each time increment (**Fig 3**), and the heatmap showing the host genes expression in response to ASFV infection (**Fig 6**), were generated using the *ComplexHeatmap* and *circlize* packages in R software. The trend curves of the viral clusters (**Fig 4**), and the graphs of the quantification of infected and activated cells (**Fig 8)** were generated using the *ggplot2* package in R software.

The schematic representations of transcriptomic and functional assays, presented in **Fig 1A** and **Fig 5A**, were made using Biorender.com under an academic license. The L60 genome representation in **Fig 4B** was made using SnapGene Viewer software (v6.2.1).

## Acknowledgments

We acknowledge Sylvain Bourgeais and Benoît Marteau from the Unité Expérimentale de Physiologie Animale de l’Orfrasière (UEPAO, INRAE) for providing access to the animal samples essential for this research. We also thank Rodrigo Guabiraba-Brito for critical reading of the manuscript.

---

Figure S1: MOI adaptation does not impact the differences observed between virulent and attenuated ASFV strains during the early phase of DNA replication.

**(A-C)** Infection rate at 24 hpi assessed by flow cytometry. PAMs from three piglet donors were plated in 12-well plates at a concentration of 1.0 x 10^6^ cells per well. 24 h later, cells were infected at the indicated MOI with virulent (L60 and OURT88/1) and attenuated (NH/P68 and OURT88/3) strains. After 24 h of infection, PAMs were stained intracellularly with a p72 antibody and infection rate was quantified using a flow cytometer. A one-way ANOVA test was performed to compare infection rate among strains conditions (*ns: non-significant*). **(B-D)** Detection of viral DNA replication by qPCR. PAMs from 3 donors were plated in 12-well plates at a concentration of 1.0 x 10^6^ cells per well. 24 h later, cells were infected at the indicated MOI with virulent (L60 and OURT88/1) and attenuated (NH/P68 and OURT88/3) strains. DNA samples of infected PAMs were collected at 2, 4, 8, 24 and 48 hpi and analyzed by qPCR for the quantification of the *B646L* gene. A two-way ANOVA test was performed to compare virulent (L60 and OURT88/1) and attenuated (NH/P68 and OURT88/3) strains replication (*\*: p<0.05*, *\*\*: p<0.01*, *\*\*\*: p<0.001*, *\*\*\*\*: p<0.0001*, *ns: non-significant*).

Figure S2: Overview of the microfluidic RT-qPCR assay results and of the global dynamics of viral genes expression.

**(A)** Number and strain-specificity of genes analyzed in the microfluidic transcriptomic assay.

**(B)** Mean number of newly expressed genes at each time post-infection for each viral strain. The threshold to differentiate expressed from non-expressed genes was settled at 25 Ct (equivalent to 35 Ct in conventional RT-qPCR since pre-amplification of cDNA lowers the Ct value by 10 in microfluidic RT-qPCR assay). A two-way ANOVA test was performed to compare the number of newly expressed genes at each timepoint between virulent (L60 and OURT88/1) and attenuated (NH/P68 and OURT88/3) strains (*ns: non-significant*, *\*\*\*: p>0.001, **: p<0.01*). **(C)** Principal Component Analyses (PCA) were performed on fold change expression data at 4, 8, 24 and 48 hpi. Each point represents a sample, colored accordingly to the viral strain used for the infection.

Figure S3: Dynamics of expression of genes identified as overexpressed at 24 hpi or with an increased transcriptional rate between 4 and 8 hpi during virulent strains infection.

**(A)** Expression profiles of the upregulated genes at 24 hpi during virulent strains infection. For each sample, the expression of each viral gene was normalized against *GAPDH* and *RPL19* expression and fold change values were normalized against the pool of all cDNA analyzed in microfluidic RT-qPCR. A two-way ANOVA test was performed to compare gene expression among times post-infection of attenuated and virulent strain infection (*ns: non-significant, *: p <0.05, **: p<0.01, ***: p<0.001, ****: p<0.0001*). **(B)** Expression profiles of the 23 genes of the cluster V3. For each sample, the expression of each viral gene was normalized against *GAPDH* and *RPL19* expression and fold change values were normalized against the pool of all cDNA samples analyzed in microfluidic RT-qPCR. A two-way ANOVA test was performed to compare gene expression among times post-infection of attenuated and virulent strains infection (*ns: non-significant, *: p <0.05, **: p<0.01, ***: p<0.001*).

**Figure S4: Profiles of expression of host genes in ASFV infected PAMs.**

**(A)** Genes specifically triggered by attenuated ASFV strain infection. The microfluidic RT-qPCR data for *IFNA1* and selected genes of host cluster H6 are shown. (**B)** Genes downregulated by both attenuated and virulent ASFV strains. Microfluidic RT-qPCR data for selected genes of host gene cluster H1 are shown. For each sample, the expression of each host gene was normalized against *GAPDH* and *RPL19* expression and fold changes value were normalized using uninfected PAMs collected at the beginning of the experiment (0 hpi). A two-way ANOVA test was performed to compare the fold change of gene expression between virulent and attenuated strain infection with non-infected condition at each time-point (*\*: p <0.05, **: p<0.01, ****: p<0.0001*).

**Figure S5: Early induction of ISGs is independent of type I interferon present in viral stocks. (A**,**B**) Detection of IFNα (**A**) and IFNβ (**B**) in viral stocks used in this study by ELISA. The threshold of detection is symbolized by the dotted line. **(C)** Induction of genes in PAMs stimulated with porcine IFNα. 5.0 x 10^5^ PAMs isolated from 3 donors were plated in 24-well plates and stimulated for 8, 16 and 24h with recombinant porcine IFNα at 1 or 250 U/mL. PRRs (*CGAS*, *ZBP1*, *EIF2AK2*), *ISG15* and *HSP70.2* induction was determined by RT-qPCR using non stimulated sample as reference. **(D)** Comparison of genes induction between NH/P68 and OURT88/3 attenuated strains. PRRs (*CGAS*, *ZBP1*, *EIF2AK2*), *ISG15* and *HSP70.2* induction after 4 and 8h of infection with NH/P68 (light blue) or OURT88/3 (dark blue) strains, detected by microfluidic RT-qPCR assay. The expression of each host gene was normalized against *GAPDH* and *RPL19* expression and fold change values were normalized using uninfected PAMs collected at the beginning of the experiment (0 hpi). **(E)** Differentially expressed genes between NH/P68 and OURT88/3 strain infection at 4, 8, 24 and 48 hpi determined by microfluidic RT-qPCR and visualized as volcano plots. Each point represents a host gene. Expression data was normalized the same way as in **(D)**. X axis represent the log_2_ of the ratio of the mean fold change of NH/P68 and OURT88/3 strains infection. Y axis represent the – log_10_ of p-value obtained by an independent t-test comparing the fold change of genotype I attenuated strains at each time post-infection. Genes with significant upregulation in NH/P68 infection are shown in light blue (Log_2_FC < 0 and p-value < 0.05), and genes with significant upregulation in OURT88/3 infection are shown in dark blue (Log_2_FC > 0 and p-value < 0.05). Non-significant genes are shown in grey.

**Figure S6: Profiles of expression of host genes in Georgia 2007/1 infected PAMs.**

For each sample, expression of each host gene was normalized against *GAPDH* and *RPL19* expression and fold change values were normalized using uninfected PAMs collected at the beginning of the experiment (0 hpi). A two-way ANOVA test was performed to compare the fold change of gene expression between genotype I virulent or attenuated strain infection and genotype II virulent Georgia 2007/1 strain infection with the non-infected condition at each time-point (*\*: p <0.05, **: p<0.01, ****: p<0.0001*).

